# Experimental and theoretical support for costs of plasticity and phenotype in a nematode cannibalistic trait

**DOI:** 10.1101/2022.02.28.482339

**Authors:** Mohannad Dardiry, Veysi Piskobulu, Ata Kalirad, Ralf J. Sommer

**Affiliations:** Max Planck Institute for Biology Tübingen; Department for Integrative Evolutionary Biology, Max Planck Ring 9, 72076 Tübingen, Germany

**Keywords:** Cost of plasticity, Cost of phenotype, cannibalism, adaptive plasticity, Markov population models

## Abstract

Developmental plasticity is the ability of a genotype to express multiple phenotypes under different environmental conditions and has been shown to facilitate the evolution of novel traits. However, while the associated cost of plasticity, *i*.*e*., the loss in fitness due to the plastic response to environment, and the cost of phenotype, *i*.*e*., the loss of fitness due to expressing a fixed phenotype across environments, have been theoretically predicted, empirically such costs remain poorly documented and little understood. Here, we use a plasticity model system, hermaphroditic nematode *Pristionchus pacificus*, to experimentally measure these costs in wild isolates under controlled laboratory conditions. *P. pacificus* can develop either a bacterial feeding or predatory mouth morph in response to different external stimuli, with natural variation of mouth-morph ratios between strains. We first demonstrated the cost of phenotype by analyzing fecundity and developmental speed in relation to mouth morphs across the *P. pacificus* phylogenetic tree. Then, we exposed *P. pacificus* strains to two distinct microbial diets that induce strain-specific mouth-form ratios. Our results indicate that the plastic strain does shoulder a cost of plasticity, *i*.*e*., the diet-induced predatory mouth morph is associated with reduced fecundity and slower developmental speed. In contrast, the non-plastic strain suffers from the cost of phenotype since its phenotype does not change to match the unfavorable bacterial diet, but shows increased fitness and higher developmental speed on the favorable diet. Furthermore, using a stage-structured population model based on empirically-derived life history parameters, we show how population structure can alleviate the cost of plasticity in *P. pacificus*. The results of the model illustrate the extent to which the costs associated with plasticity and its effect of competition depend on ecological factors. This study provides comprehensive support for the costs of plasticity and phenotype based on empirical and modeling approaches.

**Impact Summary:** A genotype able to express a range of phenotypes in response to environmental conditions, that is to demonstrate developmental plasticity, would be a Darwinian demon, able to infinitely adapt and outcompete those genotypes that require genetic change to express a phenotype fit to an environment. It has been suggested that the absence of such demons in nature is due to the cost of plasticity, *i*.*e*., developmental plasticity results in a reduction of biological fitness compared to a genotype that facultatively expresses a phenotype matching the environment. While conceptually simple, measuring the cost of plasticity in nature has proven a major challenge. We use the nematode *P. pacificus* to measure the cost of plasticity. During its development, *P. pacificus* can assume one of two possible mouth forms: predatory or non-predatory. The likelihood developing any of these two mouth forms is determined by a gene regulatory network, which itself is affected by a wide range on environmental conditions, including diet. We used two strains of *P. pacificus* and grew them on two different bacterial diets. The plastic strain was capable of switching from non-predatory to predatory mouth form depending on the diet, while the non-plastic strain could only express the predatory mouth form on either of the diets. By measuring the number eggs laid in both strain on each diet, we show that the plastic response is associated with a reduction in fecundity, thus providing a clear example of the cost of plasticity. We then use a stage-structured model to simulate the population dynamics of the plastic and the non-plastic strains. Our simulation show that the cost of plasticity is highly context dependent and its ecological ramifications can be greatly influenced by biotic and abiotic factors.

## Introduction

Changing and fluctuating environments are a hallmark of all ecosystems, affecting the life and evolution of all organisms (Sæther & Engen 2015; Pfennig 2021). The ability of an organism to respond to changing environments by expressing alternative phenotypes, *i*.*e*. phenotypic plasticity, can, in theory, facilitate adaptation, as it makes various trait optima across time and space in a given environment accessible to a genotype without the need for genetic change (Pigliucci 2001; West-Eberhard 2003; Pfennig 2021). Indeed, many case studies in plants, insects, vertebrates, and nematodes have indicated the importance of phenotypic plasticity for promoting adaptations across environments and for the evolution of novelty (West-Eberhard 2003; Moczek *et al*. 2011; Sommer 2020). We refer to this type of phenotypic plasticity that facilitates adaptation as “adaptive plasticity”. However, not all examples of phenotypic plasticity are adaptive (Murren *et al*. 2015). Additionally, plastic genotypes can vary in their degree of plasticity across different conditions (Scheiner 1993; Schlichting & Pigliucci 1998). Such assumptions imply the existence of constraints on the evolution of adaptive plasticity, and a huge and growing body of research has been dedicated to identify such hypothetical constraints (DeWitt *et al*. 1998; Callahan *et al*. 2008; Auld *et al*. 2010; Snell-Rood *et al*. 2010; Murren *et al*. 2015; Pfennig 2021). Theoretically, several factors can hinder the evolution of adaptive plasticity: limited genetic variation, weak selection, and the unreliability of environmental signals (Schlichting & Pigliucci 1998; Snell-Rood *et al*. 2010; Pfennig 2021). Most importantly, however, it has been argued that fitness costs will limit the evolution of plastic phenotypes (Murren *et al*. 2015). Such arguments are largely theoretical, and detecting the cost of adaptive plasticity remains a formidable challenge. For example, a meta-analysis of 27 studies on the cost of plasticity in animals and plants concluded that most of the studies did not have adequate statistical power to detect the cost associated with plasticity (Van Buskirk & Steiner 2009). Here, we use the plasticity model system, the nematode *Pristionchus pacificus* (Fig.1), to obtain experimental evidence for plasticity-associated costs.

**Fig. 1:**
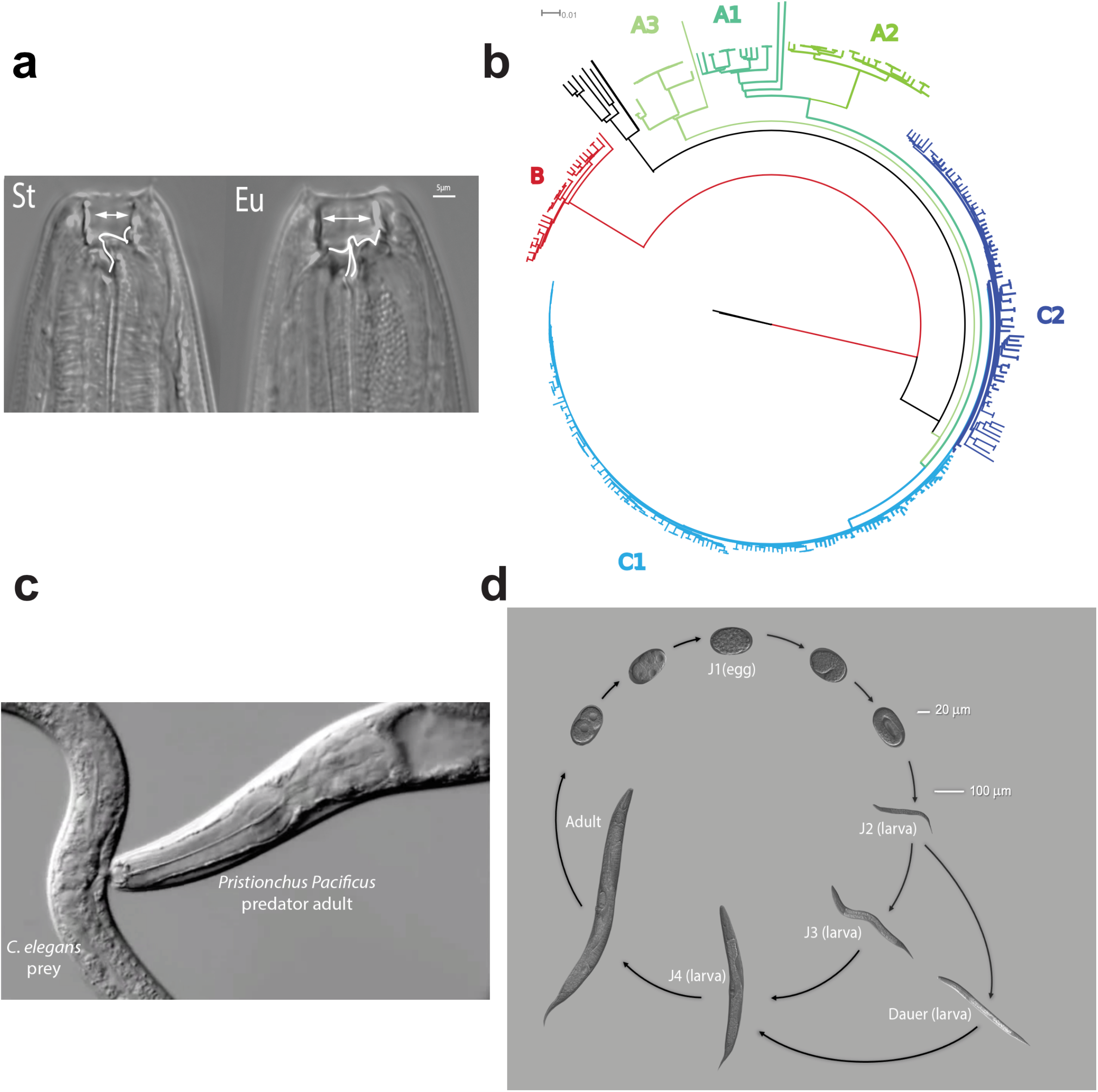
Mouth-form plasticity in *P. pacificus*: **(a)** Mouth morph dimorphism in *P. pacificus*. The bacterivorous St morph (non-predatory) possesses a single tooth with a narrow buccal cavity, whereas the predatory Eu morph (predatory) displays two teeth with a broad buccal cavity. **(b)** *P. pacificus* phylogenetic tree representing the genomic relationship between 323 *P. pacificus* wild isolates. The three major clades of *P. pacificus* are all represented in this study. Note that clades A, B, and C have approximately 1% inter-clade genomic divergence. (Adopted from Rödelsperger *et al*. 2017). **(c)** *P. pacificus* adult preying on a *C. elegans* worm. **(d)** *P. pacificus* life cycle including four juvenile stages before reaching adulthood. In stressful conditions; *e*.*g*., food depletion, juvenile worms develop into the alternatively dauer stage, which also represents the dispersal stage. Upon suitable conditions, worms exit the dauer stage and resume normal development.

To understand the costs associated with adaptive plasticity, one must consider the potential trade-offs between plastic *vs*. non-plastic genotypes in a comparative framework (Fig. 2a). Hypothetically, a non-plastic (fixed) genotype might express a mismatching phenotype in environment I. In contrast, a plastic genotype may express another phenotype more suitable for the same environment that results in higher fitness (Fig. 2a). This fitness difference between the two genotypes has been referred to as *the cost of phenotype* (Callahan *et al*. 2008; Murren *et al*. 2015). In environment II, the plastic genotype can incur lower fitness than the non-plastic genotype due to the induction of a mismatching phenotype (Fig. 2a). This hypothetical fitness trade-off associated with the plastic genotype has been referred to as *the cost of plasticity* (Callahan *et al*. 2008; Murren *et al*. 2015). The *cost of phenotype* can be described as the organismal fitness reduction due to devoting resources to the continuous production of resource-demanding phenotypes. In contrast, *the cost of plasticity* can be defined as the price paid in fitness by a highly-plastic genotype compared to a less plastic one (Callahan *et al*. 2008; Murren *et al*. 2015). Understandably, the terms ‘cost of phenotype’ and ‘costs of plasticity’ are, by virtue of their definitions, ripe for confusion (Auld *et al*. 2010; Pfennig 2021), and they can be only applied within a comparative framework (Callahan *et al*. 2008). A comprehensive analysis of these constraints would adequately improve our understanding of the role of plasticity in adaptive evolution. However, empirical studies on the costs of phenotype and plasticity remain scarce, especially in metazoans. This has been largely due to two reasons: Firstly, in the wild, conditions often cannot be properly controlled, nor can the effects of various factors be delineated. Secondly, laboratory experiments are time consuming and large organisms cannot be easily investigated. To study the constraints of plasticity, we make use of natural isolates of nematodes that, given their small size and rapid reproduction, can be examined under laboratory conditions.

**Fig. 2:**
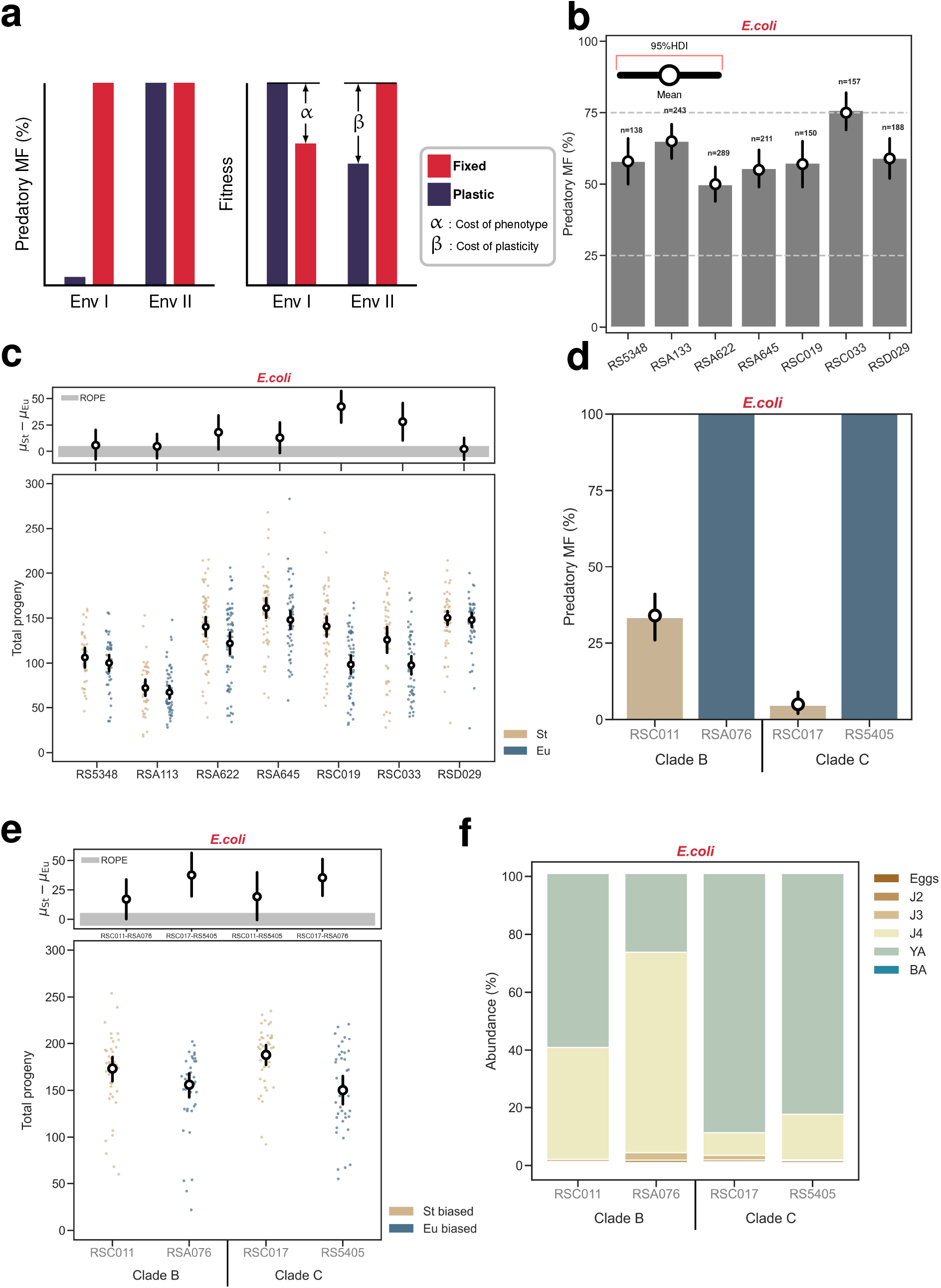
Within and between strains costs of phenotype in *P. pacificus*: **(a)** A hypothetical scenario illustrating the cost of plasticity and the cost of phenotype: a plastic genotype switches from the non-predatory to the predatory mouth form when grown in environment II. In contrast, a non-plastic (fixed) genotype constantly expresses the predatory mouth form regardless of environmental changes. Costs of phenotype and plasticity are, by definition, exclusively meaningful in comparative studies in environments that can be described as adaptive and non-adaptive with regards to a given trait. **(b)** Percentage of predatory mouth form of seven intermediate *P. pacificus* genotypes used in this study. The number of worms assayed for each strain (n) is indicated above each of the 95% highest density interval (HDI) for the means. **(c)** Overall fecundity of the same seven intermediate strains. On average between 51 and 47 predatory and non-predatory mothers per strain were scored, respectively. Two comparisons, RSC019 and RSC033, are outside the region of practical equivalence (ROPE), *i*.*e*., means in theses comparisons are different; two other comparisons, RSA622 and RSA645, partially overlap with the ROPE, and the rest of the comparisons include the ROPE, implying no difference. **(d)** Percentage of predatory mouth form of four biased *P. pacificus* wild isolates representing clades B and C, respectively. For each strain, three biological replicates with a total number of 150 worms per strain were scored. **(e)** Overall fecundity of the same biased strains. Both non-predatory-biased strains showed credibly higher overall fecundity than the predatory-biased strains. **(f)** Developmental speed for the biased strains from the two clades. For RSC011, 263; RSA076, 232; RSC017, 320; and RS5405, 306 individuals were staged. Worms were staged according to the following developmental stages: E= eggs; J2, J3, J4 = juvenile stages; YA= young adults with no eggs inside the uterus; BA= breeding adults with eggs inside the uterus. In (b) and (d), the 95%HDI for each strain was estimated using a Bayesian approach to estimate the probability of expressing the predatory mouth morph based on the observations; the 95% HDI for the means and the difference in means in (c) and (e) was calculated using Kruschke’s BEST method. We used [-5,5] interval as our ROPE, *i*.*e*., differences of means within this interval are practically equal to no difference; the same ROPE and was used for all the analyses in this manuscript (see *Supplementary Methods*).

The nematode *P. pacificus* is an established model system for studying phenotypic plasticity (Sommer & McGaughran 2013; Sommer *et al*. 2017). The developmentally plastic mouth of *P. pacificus* can exhibit two distinct forms; the eurystomatous (predatory) morph with a wide stoma and hooked-like teeth, or the stenostomatous (non-predatory) morph with a narrow stoma and a single tooth (Fig.1a) (Bento *et al*. 2010). *P. pacificus* is a hermaphroditic nematode and the use of isogenic cultures has facilitated the elucidation of genetic and epigenetic mechanisms underlying this irreversible switch. Specifically, the sulfatase-encoding *eud-1* gene was identified as the key developmental switch that is regulated by various environmental factors and epigenetic mechanisms, and directs a downstream gene regulatory network consisting of more than 20 identified proteins including structural components of mouth formation (Ragsdale *et al*. 2013; Kieninger *et al*. 2016; Bui *et al*. 2018; Namdeo *et al*. 2018; Sieriebriennikov *et al*. 2018; Sieriebriennikov *et al*. 2020; Sun *et al*. 2022). Importantly, worms respond to surrounding environmental cues to adopt their mouth form in a strain-specific manner and various environmental stimuli, including temperature, culturing condition, crowding and diet have been shown to influence mouth-morph ratios (Werner *et al*. 2017; Werner *et al*. 2018; Lenuzzi *et al*. 2021). Principally, three major features assist in studying *P. pacificus* mouth-form plasticity. First, the vast collection of naturally occurring wild isolates with hundreds of *P. pacificus* strains being sequenced, accordingly resulted in a highly resolved phylogeny of diverse populations (Fig.1b) (Rödelsperger *et al*. 2017). Interestingly, culturing these isolates on the laboratory bacterium *E. coli* displays a range of mouth-morph ratios, while some express intermediate mouth-form ratios (see below) (Ragsdale *et al*. 2013). While the parallel formation of both mouth forms under the same environmental condition represents an unusual type of plasticity that is not seen in the majority of plastic traits, it has allowed unprecedented insight into associated molecular mechanisms and the identification of a large gene regulatory network (Ragsdale *et al*. 2013; Kieninger *et al*. 2016; Bui *et al*. 2018; Namdeo *et al*. 2018; Sieriebriennikov *et al*. 2018; Sieriebriennikov *et al*. 2020; Sun *et al*. 2022). Second, morphological mouth-form plasticity is coupled to behavioral plasticity. Specifically, the predatory form enables predation and cannibalism on other nematodes, while such animals can still feed on bacteria. In contrast, the non-predatory form obligates worms to feed on bacteria (Fig.1c) (Wilecki *et al*. 2015). This extension of morphological plasticity to behavior is thought to eliminate resource competitors via predation and the expansion of nutrition (Quach & Chalasani 2020). Finally, *P. pacificus* is a thoroughly-studied soil nematode that is reliably found in association with scarab beetles with recent studies describing the dynamics and succession of nematodes on the beetle carcass after the insect’
ss death (Fig.1d) (Renahan *et al*. 2021).

Switching between the predatory and non-predatory mouth forms is a specific example of phenotypic plasticity, in which an irreversible decision that occurs during the development of *P. pacificus* via a bi-stable developmental genetic switch (Sieriebriennikov *et al*. 2018). The bias of the developmental switch determines the ratio of mouth morphs with substantial natural variation between populations of *P. pacificus* (Ragsdale *et al*. 2013). In this respect, mouth-form plasticity in *P. pacificus* is more akin to the switch between lytic and lysogenic cycles in bacteriophage *λ* (Ptashne 1986) than wing pattern polyphenism in butterflies that is seasonally controlled (Nijhout 1994). Importantly, the relative simplicity of mouth-morph plasticity in *P. pacificus*, as well as its isogenic husbandry of genetically diverse strains, makes it ideal to study different facets of phenotypic plasticity. Here, we took advantage of these features to perform a systematic analysis of mouth morphs and their associated costs and extended our empirical findings via simulating ecologically relevant scenarios in spatially-homogeneous and spatially-structured populations using empirically-derived life history parameters.

## Methods

Please see the Supporting Information.

## Results

### Within and between strains comparisons reveal a cost of phenotype

To measure the cost of phenotype, we took advantage of the extensive collection of *P. pacificus* natural isolates and selected seven strains with intermediate mouth-morph ratios from across the *P. pacificus* phylogeny (Fig.1b) (Rödelsperger *et al*. 2017; Lightfoot *et al*. 2021). The ratios of these strains were considered intermediate since they were neither predominantly predatory nor predominantly non-predatory on the standard laboratory food source *E. coli* (Fig. 2b, *SI Appendix*; Table. S4). It should be noted that “intermediate” in this context does not imply a 50-50 chance of expressing either of the mouth morphs, but merely indicates that both alternative morphs can be easily found in the lab. These strains allow testing whether there is a cost of phenotype within the same genetic background. Note that such strains are rare in nature and were obtained only because of the large available collection of *P. pacificus* wild isolates.

To measure the cost of phenotype, we selected overall individual fecundity as primary fitness parameter to capture reproductive capacity of *P. pacificus* hermaphrodites via selfing (Haldane 1937; Orr 2009). Testing for daily fecundity showed that the majority of progeny were laid within a window of 62 hours after maturation (nearly 91%) in an overall window of appr. 158 hours of total egg-laying (*SI Appendix*; Fig. S1a& Table. S2). The number of eggs laid in this window provide a reasonable estimate of the life-time fecundity. In these intra-genotype comparisons, we found a tendency in non-predatory animals to have more progeny than predatory worms. Specifically, the estimated differences in the mean value of fecundity between non-predatory and predatory animals based on the data showed strong and/or partial support in four out of seven comparisons (Fig. 2c, *SI Appendix*; Table S1; Table S6). These findings suggest that the production of the predatory mouth morph can incur a fitness cost. This observation is in concert with a previously report on the slower rate of development in nematodes exhibiting the predatory morph in comparison to non-predatory worms (Serobyan *et al*. 2013).

Afterwards, we measured fecundity and developmental speed in *P. pacificus* natural isolates that show a biased mouth-morph ratio, *i*.*e*., strains that would produce an abundance of non-predatory or predatory mouth morphs on the standard laboratory food source *E. coli* (Fig. 2d, *SI Appendix*; Table S4). We selected two pairs of closely-related strains from the diverged clades B and C of *P. pacificus* from La Réunion island (Rödelsperger *et al*. 2014). We found that in both pairs, the non-predatory-biased strains produce more overall progeny than the predatory-biased strains (Fig. 2e, *SI Appendix*; Table S1; Table S7). Specifically, giving the same time window of the first 62 hours, the St-biased strains had a 21% and 17% higher fecundity in clades B and C, respectively (*SI Appendix*; Fig. S2b& Table. S2). Similarly, the non-predatory-biased strains showed a higher developmental speed (Fig. 2f, *SI Appendix*; Table. S3). For example, 75 hours after egg-laying, nearly 60% of the non-predatory-biased strain RSC011 reached adulthood; whereas only 27% of the predatory-biased strain RSA076 reached the same stage. Note that inter-clade comparisons show considerable differences in these isolates’ developmental speed, which is due to the genetic background. Together, both, our pairwise comparisons of total eggs laid by predatory and non-predatory individuals in intermediate strains, and the pairwise comparisons of fecundity and developmental speed in four biased strains from two different *P. pacificus* clades clearly illustrate the cost of producing the predatory phenotype.

### Across-conditions testing indicates a cost of plasticity

Next, we wanted to determine if a cost of mouth-form plasticity exists in *P. pacificus*. Such a cost of plasticity would be eminent when testing a non-plastic genotype relative to a plastic genotype under different conditions (DeWitt *et al*. 1998; Pigliucci 2001; Callahan *et al*. 2008; Murren *et al*. 2015). Therefore, we performed a cross condition test by conducting experiments on two distinct food sources, the standard *E. coli* condition used in the previous section, and a *Novosphingobium* diet. The bacterial species *Novosphingobium* was found to be naturally associated with *P. pacificus* and was proven to increase intraguild predation in the *P. pacificus* reference strain PS312 (Akduman *et al*. 2018; Akduman *et al*. 2020). However, this association was never studied in non-domesticated wild isolates of *P. pacificus*. Therefore, we grew two of the biased strains with different mouth-morph ratios on *Novosphingobium;* the highly non-predatory-biased strain RSC017 and the highly predatory-biased strain RS5405. Indeed, RSC017 showed a substantial increase of the predatory morph of 84% on *Novosphingobium*, indicating strong plasticity. In contrast, the predatory-biased strain RS5405 remained highly predatory in the new condition (Fig. 3a, *SI Appendix;* Table. S4). Thus, we established two distinct food conditions that differentially affect plasticity levels of the two isolates. Henceforth, we refer to RSC017 and RS5405 as plastic and non-plastic strains, respectively.

**Fig. 3:**
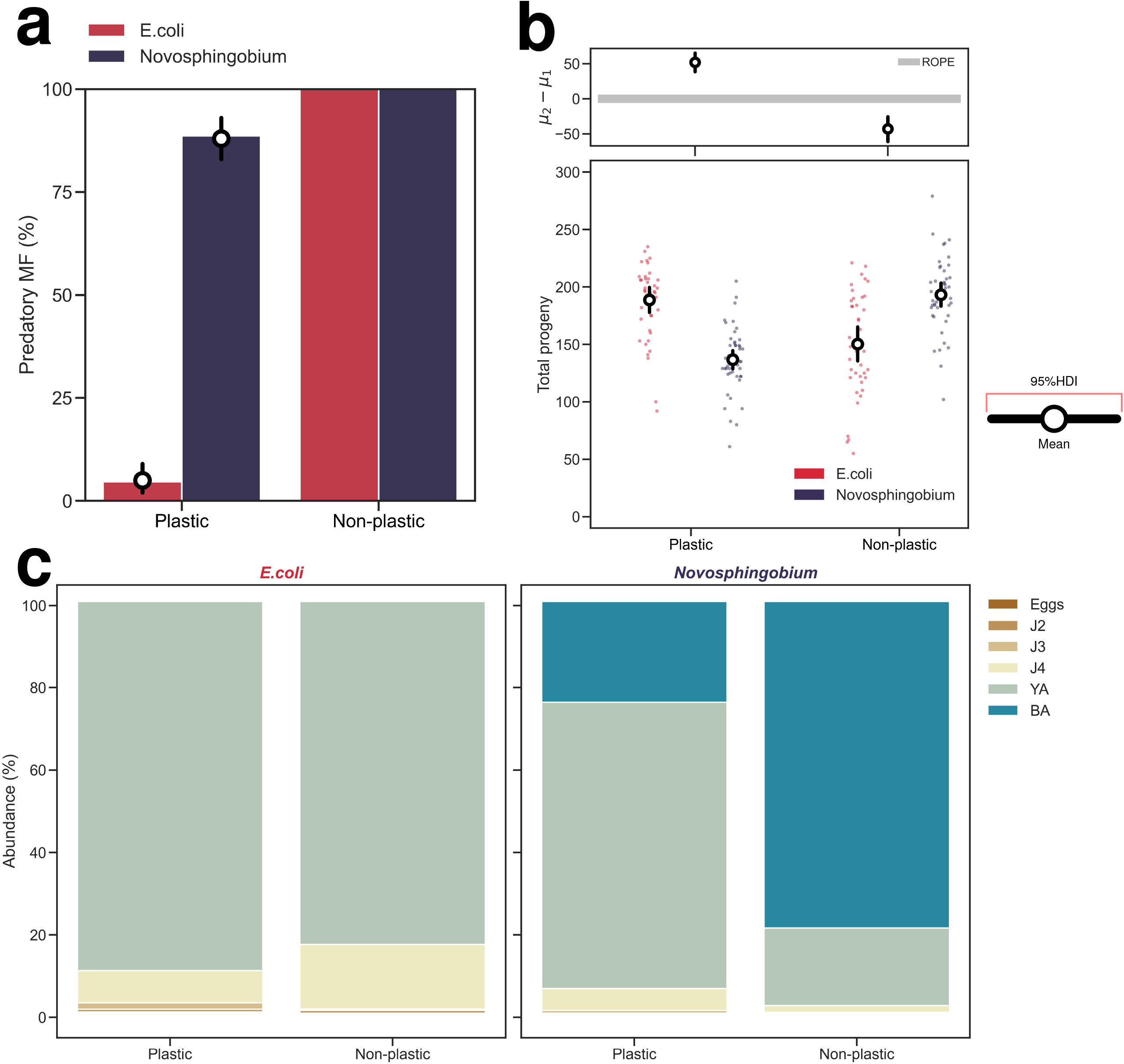
Cost of plasticity across conditions: **(a)** Percentage of predatory mouth form on *E. coli* and *Novosphingobium* for *P. pacificus* strains, representing both biased mouth-morph ratios. A total of 150 animals were used to score the mouth-morph ratio for each strain per condition, indicating three biological replicates. **(b)** Overall fecundity of the *P. pacificus* strains on both food conditions. The top panel indicates the 95% HDI for the estimated difference in means for each pairwise comparison calculated using Kruschke’s BEST method (see *Supplementary Methods*). Comparable numbers of mothers were used: the plastic strain (*E. coli*=40, *Novosphingobium*=47); the non-plastic strain (E. *coli*=40, *Novosphingobium*=43) **(c)**. Developmental speed for the two genotypes on both conditions. Individual worms were staged 75 hours after mothers were killed; for the plastic strain (*E. coli*=320, *Novosphingobium*=301); the non-plastic strain (*E. coli*=306, *Novosphingobium*=489) individuals were staged. Worms were staged according to the following developmental stages: E= eggs; J2, J3, J4 = juvenile stages; YA= young adults with no eggs inside the uterus; BA= breeding adults with eggs inside the uterus.

Theoretically, the cost of plasticity would be displayed in the strain that exhibits a change in mouth-morph ratio upon altering food conditions. Accordingly, we would expect to detect the highest effect on fitness in the plastic strain, and vice versa. Indeed, we found that the plastic strain has lower fecundity and slower developmental speed on *Novosphingobium* when compared to the non-plastic strain (Fig. 3b-c, *SI Appendix*; Table. S1,3, Table S8). Thus, a strain that plastically responds to a dietary change with the formation of the predatory mouth morph exhibits reduced fitness under these novel conditions indicating a cost of plasticity. In contrast, the non-plastic strain exhibits higher levels of fecundity and developmental speed on *Novosphingobium* (Fig. 3b-c, *SI Appendix*; Table. S1,3). Thus, a strain that is preferentially predatory under both food conditions exhibits increased fitness when exposed to this new diet. Taken together, these findings indicate a cost of mouth-morph plasticity in response to dietary induction. These observations raise a fascinating question: which cost plays a larger role in shaping the population dynamics and, consequently, the evolution of mouth-morph ratios?

### The cost of phenotype maximizes the benefits of plasticity

To investigate how the cost of plasticity and the cost of phenotype would manifest in the wild, we constructed a stage-classified model to simulate population dynamics of the plastic and the non-plastic strains on both tested food sources (Fig. 4a). For modeling, we used the fecundity measurements from the lab and scaled the developmental rates of the model based on the laboratory estimates of developmental speed of *P. pacificus* (see *Supplementary Methods*). First, we tested population dynamics of the selected strains in separation, *i*.*e*., without interactions or competition. Surprisingly, the change from *E. coli* to *Novosphingobium* has only a minor effect on the final population size of the plastic strain (Fig. 4b). The reduction in fecundity on *Novosphingobium* relative to *E. coli* is presumably compensated by the increase in developmental speed on *Novosphingobium*. To test the hypothesis that faster developmental speed was indeed compensating for the cost of plasticity, (*i*.*e*., lower fecundity), we simulated the dynamics of the plastic stain by assuming no change in developmental speed. The results of this simulation confirmed this expectation (Fig S6). In contrast, in the non-plastic strain the increase in fecundity and developmental speed on *Novosphingobium* results in a higher frequency of all developmental stages compared to its dynamic on *E. coli* (Fig. 4c). Importantly, the between strains cost of phenotype is clearly displayed when comparing the frequencies of the two strains on *E. coli* (Fig. 4b-c). Thus, comparing both populations’ trajectories without involving interactions reveals that the cost of phenotype has a larger effect on the population dynamics than the cost of plasticity.

**Fig. 4:**
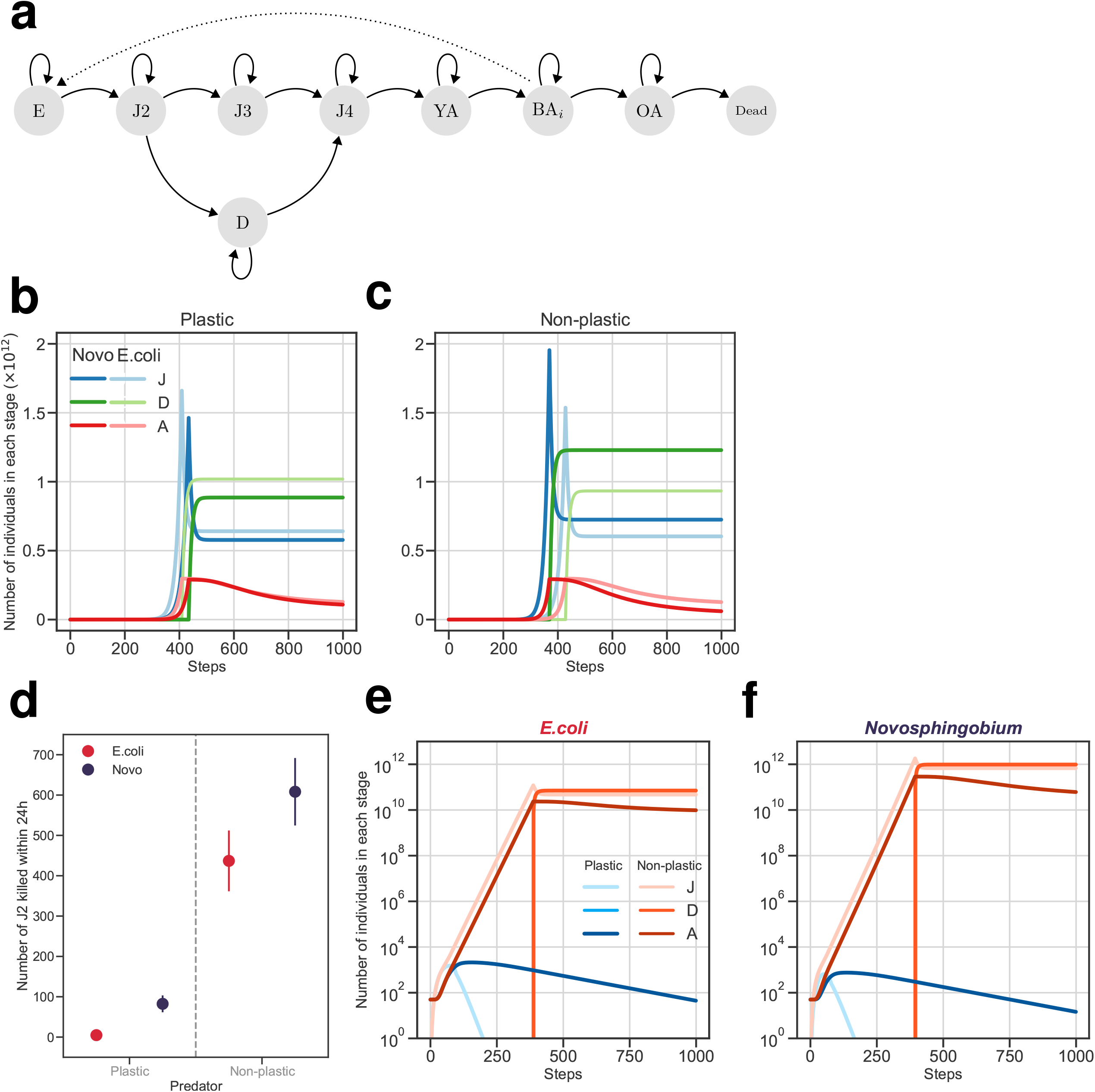
Costs of phenotype and plasticity in a spatially-homogeneous population: **(a)** Life cycle of *P. pacificus* as a Markov chain (E: egg, J2-4: juvenile stages, D: dauer, YA: young adult, BA_*i*_: breeding adult of the day *i*, OA: old adult). Note that J1 larvae in *P. pacificus* remain in the egg shell and are considered part of E in our model. Solid arrows represent the transition between different developmental stages. Egg-laying by BA adults is indicated by a dotted arrow. Five different breeding adults are included in the model (BA_1_ to BA_5_). **(b-c)** Population dynamics of the plastic and the non-plastic strains on *E. coli* and *Novosphingobium*. Note that the end point of 1000 steps represent an arbitrary endpoint, which roughly corresponds to 10 generations. Food will be gone long before this end point, as evident by the production of dauer larvae. Simulations based on empirical data demonstrate the differential response of the strains on both food conditions. The number of adults and dauer larvae for the plastic strain in *E. coli* relative to their counts on *Novosphingobium* are 1.05 and 1.09, respectively. For the non-plastic strain, the number of adults and dauer larvae in *E. coli* relative to their counts on *Novosphingobium* are 2.28 and 0.82, reflecting the faster YA to BA_1_ and higher fecundity of the non-plastic strain on *Novosphingobium*. Note that J is the sum of all juvenile stages. Upon the depletion of food, juvenile stages and eggs stop transitioning into the next developmental stage, except for J2, which develops into dauer larvae. **(d)** Standard inter-strain corpse assay. Each dot represents a mean of five replicates, and error bars represent standard deviation. For each replicate, 20 adult predators were added to ≈ 3000 J2 prey, and corpses were screened after 24 hours. **(e-f)** Simulation of the effect of with-strain predation on population dynamics in a spatially-homogeneous population. Using predation rate estimates from the corpse assay, we simulated the interaction of the plastic and the non-plastic strain on *E. coli* and *Novosphingobium*. In both conditions, the non-plastic strain the non-plastic strain drives the plastic strain into extinction. Simulations in (b), (c) start with 50 YAs of a strain. The initial food supply, *S*_0_ = 10^12^. On E. coli, for both strains, *γ*_E_ = 0.0415, *γ*_J2_ = 0.055, *γ*_J3_ = 0.085, *γ*_J4_ = 0.07, *γ*_YA_ = 0.1, *γ*_Bi_ = 0.0415, *σ*_OA_ = 0.995 (see *Supplementary Methods*). On *Novosphingobium, γ*_YA_ = 0.13 for the plastic strain and *γ*_YA_ = 0.4 for the non-plastic strain to account for the change in the developmental speed observed in the experiment. The same transition probabilities and survival are used for the rest of the simulations. Simulations in (e) and (f) start with 50 YAs of each strain. Predation rates: on *E. coli, η*_RSC017_ = 1.7 × 10−4 and *η*_RS5405_ = 3.3 × 10−4; on *Novosphingobium, η*_RSC017_ = 6.4 × 10−5 and *η*_RS5405_ = 4.7 × 10−4. The same predation rates are used in the subsequent simulations.

### The cost of plasticity manifests in a competition setup

In nature, *P. pacificus* does not occur in isolation, rather it competes with other nematodes over resources. Additionally, given the coupling between morphological and behavioral plasticity, predatory worms are able to predate while non-predatory worms are not. Testing the costs of plasticity and phenotype in a competition setup might shed light on the evolution of the predatory mouth morph. Therefore, we first tested if predation rate positively correlates with the proportion of predatory individuals in wild isolates. To avoid the compounding effect of relatedness on predation (Renahan & Sommer 2021), we selected *C. elegans* as prey for *P. pacificus* predators. Indeed, testing nine *P. pacificus* wild isolates with different mouth morph bias, shows that morphological and behavioral plasticity positively correlate (*SI Appendix*; Fig. S3). Second, we measured predation rates of the plastic and the non-plastic strains against one another by testing predation rates over the two food sources *E. coli* and *Novosphingobium* (Fig. 4d).

Next, we used the experimentally obtained predation values for each food source to simulate the effect of interactions between strains on their dynamics in a spatially-homogeneous population (see *Supplementary Methods, SI Appendix*; Fig. S4). Specifically, we used these estimates to simulate the interactions between the two isolates in a population with an equal number young adults form the plastic and the non-plastic strains at the start of the simulation. Notably, simulated populations were completely dominated by the non-plastic strain for both food conditions. In addition, rapid elimination of the plastic strains prevents the formation of its dauer larvae, as J2 animals of this strain were completely eradicated by the non-plastic strain (Fig. 4e, f). Thus, the cost of plasticity greatly affects the dynamics of the plastic strain in a spatially-homogenous population.

### Spatial structure significantly affects population dynamics

While modeling the interaction of the plastic and the non-plastic strains in a population without any spatial structure is informative, a more realistic scenario would involve dispersal from different populations upon the depletion of food on the beetle carcass, and competition over the nutrient-rich carcasses in the vicinity. Exploring such scenarios in the lab would be a tremendous undertaking. Therefore, we extended our model to include a stepping-stone migration scenario to illustrate the effect of costs of plasticity and phenotype on the competitive dynamics between *P. pacificus* strains in a structured population. We constructed a simple structured population by arranging n localities in one dimension. Each simulation starts with 50 young adults (YAs) of the plastic strain in the first locality and 50 YAs of the non-plastic strain in the n^th^ locality, with rest of localities being empty. All the localities contain a fixed amount of resource and dauer larvae migrate with a fixed rate from a food-poor locality to a neighboring food-rich locality (Fig. 5a). The simulation concludes when all the food in every locality has been depleted.

**Fig. 5:**
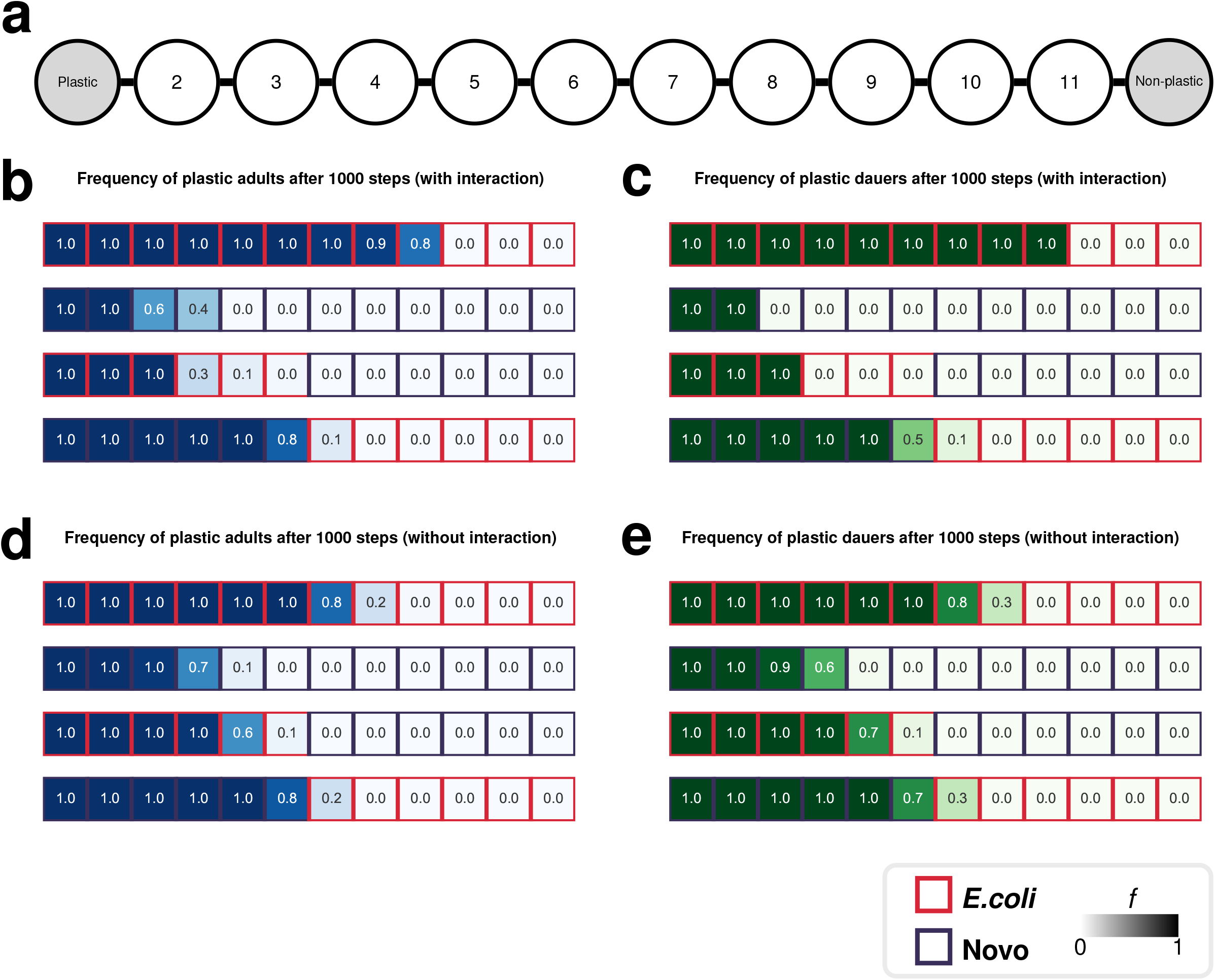
Costs of phenotype and plasticity in a spatially-structured population: **(a)** A structured population, consisting of 12 localities are arranged in a line. Each simulation starts with 50 YAs of the plastic strain on the first locality and 50 YAs of the non-plastic strain on the 12^th^ locality. At each step, *ω* dauer larvae migrate from population *i* to *j* if *j* has more food than *i*. **(b-c)** The frequency of the plastic strain adults (YA, BA_*i*_, OA) (b) and dauer larvae (c) across 12 localities with interaction (i.e., predation) after 1000 steps. As previously noted, the end point of 1000 steps represent an arbitrary endpoint, which roughly corresponds to 10 generations. Food will be gone long before as evident by the production of dauer larvae. **(d-e)** The frequency of the plastic strain adults (YA, BA_*i*_, OA) (d) and dauer larvae (e) across 12 localities assuming no interaction between the two strains. In this scenario, the frequency of the plastic strain in the metapopulation is a function of developmental speed and fecundity only. At the start of the simulation, for each strain in localities 1 and 12, *n*_E_ = *n*_J2_ = *n*_J3_ = *n*_J4_ = *n*_BA_ = *n*_OA_ = 0, and *n*_YA_ = 50. while *m* = 0.1. The initial food supply, S_0_ = 10^12^ in the entire metapopulation.

Based on these simulations, on *E. coli*, higher fecundity of the plastic strain allows adults of this isolate to completely dominate the structured population even in the face of predation (Fig. 5b, *SI Appendix*; Fig. S5a). Although the predation rate of the non-plastic strain is higher, even dauer larvae of the plastic strain continues to fully dominate the structured population (Fig. 5c). These results support a considerable cost of phenotype for the non-plastic strain in the spatially-structured population. In addition, in a scenario without predatory interactions, the frequency of the plastic strain decreases only marginally (Fig 5b *vs*. d, c *vs*. e). This finding results from the change in the number of migratory dauer larvae of the non-plastic strain (*SI Appendix*; Fig. S5a,b). Most importantly, the pace in which the plastic population grows, results in exceptionally high numbers of predators belonging to the plastic strain, which outcompete the non-plastic strain in the presence of interaction. Thus, the cost of phenotype substantially influences the abundance of the non-plastic strain, in particular in the presence of interactions.

On *Novosphingobium*, higher fecundity and faster developmental speed of the non-plastic strain turns this isolate into a formidable adversary for the plastic strain. Therefore, the frequencies of adults and dauer larvae of the plastic strain are extremely reduced in the structured population (Fig. 5b, c). However, when interactions are limited, in contrast to *E. coli*, the frequencies would slightly increase (Fig 5b *vs*. d, c *vs*. e). This is due to the non-plastic strain profiting from a higher growth rate and higher predation on *Novosphingobium*, but only higher growth when interactions are eliminated (*SI Appendix*; Fig. S5c,d). Thus, the cost of plasticity would greatly affect the abundance of the plastic strain when competing with a predator under this condition.

### Initial food source also affects population dynamics

To capture how significantly the costs of plasticity and phenotype would affect the dynamics of structured populations, we simulated two scenarios where each isolate would start with a favorable food source; *E. coli* for the plastic strain, and *Novosphingobium* for the non-plastic strain, or the unfavorable food source; *Novosphingobium* for the plastic strain, and *E. coli* for the non-plastic strain (Fig. 5b-e). A pair of food sources were labeled “favorable” or “unfavorable” for a strain given the relative fecundity of the strain on each source. Interestingly, the results indicate that the initial condition in which each population starts dramatically affects which strain would ultimately dominate the structured population. When the conditions are favorable for both strains, the cost of plasticity of the plastic strain is greater than the cost of phenotype of the non-plastic strain. In contrast, the relationship between the costs reverses under conditions that are unfavorable to both strains. Thus, the interaction of the cost of phenotype and the cost of plasticity is context dependent. Together, these simulations reveal that spatial structure and initial food sources could affect the population dynamics with different consequences for the costs of plasticity and phenotypes on the two isolates. However, such projections about the population dynamics of these strains of *P. pacificus* should be taken with caution, as many aspects of *P. pacificus* population dynamics and its dispersal patterns in the wild remain poorly understood.

## Discussion

Experiential detection of the costs associated with plasticity, especially in metazoans, has proved to be a daunting challenge. For instance, the predator-induced spine of *Daphnia pulex* was reported to show mild support for both the costs of production and maintenance (Scheiner & Berrigan 1998). Similarly, in the Scandinavian frog, *Rana temporaria*, the costs of metamorphic size were shown to exhibit a plasticity cost in southern populations, whereas northern populations displayed no such costs (Merilä *et al*. 2004). Van Buskirk and Steiner (2009), in their meta-analysis concluded that costs of plasticity are mostly low, if existing at all. However, the same authors suggested that these costs may influence adaptive evolution under stressful conditions. Additionally, meta-analysis on aquatic gastropods argued for further empirical investigations to better quantify the energetic costs of plasticity of shell formation (Bourdeau *et al*. 2015). A more recent study on the cannibalistic cane toads, signifies favoring canalized defenses over plasticity, providing the high cost of plasticity rather than the cost of phenotype (Devore *et al*. 2021). Together, this diversity of findings indicates the need for establishing a comprehensive empirical framework to address both theoretical and conceptual asserts.

The results obtained in *P. pacificus*, likely benefited from the binary and easily distinguishable state of the polyphenic trait and the isogeneic nature of all tested strains, which facilitate empirical measurements of fitness components, *i*.*e*., fecundity and developmental speed, in the laboratory. Additionally, accounting for two fitness components assisted in the transition from abstract measures to simulating a range of ecologically relevant scenarios. Our study complements previous knowledge with a systematic analysis of defined costs in an evolutionary adaptive trait.

It has been argued that resource polyphenism - *i*.*e*., the environmental induction of alternative phenotypes to use different resources, such as the development of cannibalistic morphs as a response to environmental stress (Pfennig & McGee 2010) – is the most relevant of discrete plastic response. Cannibalism provides trophic and survival advantages by either extending energy resources or eliminating competition (Church & Sherratt 1996; Claessen *et al*. 2004). It has been suggested that the predatory mouth form in *P. pacificus* boosts survivorship under severe conditions (Serobyan *et al*. 2014), and reduces competition on the basis of genomic relatedness (Lightfoot *et al*. 2021). Nevertheless, various *P. pacificus* natural isolates are either predominantly non-predatory or intermediately so. Our results suggest that, in isolation, the fitness payoff incurred by the predatory-biased population makes it inferior to the non-predatory-biased strain (Fig. 4b,c). Strikingly, our computational model indicates this cost of phenotype to be more detrimental when both isolates are interacting in a spatially-homogenous population. The effect of growth rate, developmental speed, and predation are highly context dependent, as shown by our simulations under different starting conditions, resulting in different population dynamics (*SI Appendix*; Fig. S5).

The effect of population structure and the non-homogenous distribution of resources in the environment on the outcome of competition between a plastic and non-plastic strain illustrates the complex nature of the ecological consequences of the cost of plasticity. The role of phenotypic plasticity, and dispersal, in invasions have long been appreciated (Sharma *et al*. 2005), based on the assumption that plasticity provides a “Jack-of-all-trades” strategy. This assumption has been challenged (Hulme 2007), and does not explain the pattern we observe in our model. The models proposed to predict the population-level consequences of plasticity thus far (reviewed in Wennersten and Forsman (2012)) have been almost entirely conceptual. A promising recent attempt by Brass *et al*. (2021) incorporates plasticity in a continuous-time stage-structured model to predict the ecological effects of plasticity, but their model differs from ours, since they include plasticity as maternally-determined phenotypic variation within a species and do not explore the effect of population structure nor the non-homogenous resource distribution on the cost of plasticity. The effects of spatial heterogeneity and dispersal on the evolution of plasticity have been explored before, e.g., Scheiner and Holt (2012) and Edelaar *et al*. (2017), but our results are not comparable to those, since our model lacks any evolutionary component, e.g., mutation, recombination, etc., and is solely concerned with the short-term ecological consequences of plasticity in *P. pacificus*.

Taken together, our results suggest a four-pronged explanatory framework, combining the *cost of plasticity, cost of phenotype*, environmental influence, and population structure, each playing a crucial role in adaptive plasticity. However, several questions remain to be answered. For example, measuring predation dynamics and migration rates on beetle carcasses can increase the accuracy of modeling approaches. Also, predator consumption might differ as a functional response to prey density, given search, handling time, foraging efficiency, and predation risks (Solomon 1949; Holling 1959a, b; Lima *et al*. 1985; Sentis *et al*. 2013). Additionally, in nature, nematode mobility is not restricted to a one-dimensional dispersal. Thus, such parameters merits further empirical and theoretical analyses. Finally, a key question that was hardly identified in other plastic systems is the molecular machinery underlying the production and maintenance of plasticity (Pigliucci 2001; Murren *et al*. 2015). In *P. pacificus*, the readily available molecular techniques permit such potential investigations. In conclusion, this study integrates empirical and theoretical approaches to emphasize how different types of costs influence the evolution of adaptive plasticity, while setting the stage for further investigations.

## Supporting information

Supplementary Mehtods

## Data availability

The software used to run all simulations and conduct all the data analysis was written in Python 3.10.4. For reproducibility, the code and the raw experimental data are available at (https://github.com/Kalirad/cost_of_plasticity).

## Acknowledgments

We thank Dr. Matthias Herrmann and Metta Riebesell for *P. pacificus* life cycle image; Dr. James Lightfoot for the predation image; the La Réunion field team for strains isolation; Drs. Kohta Yoshida and Christian Rödelsperger, and all members of the Sommer lab for discussions.

## Notes

### Competing Interest Statement

The authors have declared no competing interest.

### Summary of Updates

The main text has been revised and the figures updated.

