## Supplementary Mehtods for "Experimental and theoretical support for costs of plasticity and phenotype in a nematode cannibalistic trait"

Supplementary Methods

#### Details of the laboratory assays

### **Bacterial & nematodes strain culture and maintenance**

The two bacteria used as food source under monoxenic conditions were the standard *E. coli* lab strain OP50 and the naturally *Pristionchus-*associated *Novosphingobium* sp. L76 (Akduman *et al.* 2018). OP50 was grown overnight at 37°C in Lysogeny broth medium (LB) without shaking, while *Novosphingobium* sp. L76 was grown overnight at 30°C in Lysogeny broth medium (LB) at 157 rpm. On the following day, 6-cm nematode growth medium (NGM) Petri-dishes were seeded with 300µl of *E. coli* OP50 or *Novosphingobium* and left for overnight incubation (Sieriebriennikov *et al.* 2020). Nematodes were reared on the seeded NGM plates at 20°C. Three adults were passed to new plates every 5 days for *E. coli*, and every 4 days for *Novosphingobium*; giving the difference in developmental speed.

### **Mouth form phenotyping**

Mouth-form scoring was performed on a ZEISS SteREO Discovery.V20 microscope, PlanApo S 1.5x objective with eyepiece PL 10x/23 Br.foc. Mouth-form phenotype was identified according to the mouth width and the shape of the dorsal tooth of young adults as previously reported (Bento *et al.* 2010). For all experiments, three replicates were scored at 20°C on 300µl of the relevant food. The total number of worms scored per strain is as follows: Intra-strain analysis (RS5348= 138, RS113= 243, RSA662= 289, RSA645= 211, RSC019= 150, RSC033= 157, RSD029= 188). In all other analyses, i.e., inter-strain analysis, plasticity cost, and predation assays, we used 150 animals per strain.

### **Overall and daily self-fecundity measure**

Maintenance cultures were first bleached to obtain synchronized eggs before starting an experiment. Bleaching protocol was performed as previously reported in (Stiernagle). Upon synchronization, J4 larvae were individually isolated on separate plates spotted with 20µl of the relevant bacteria. The next day, when worms are young adults, the mouth form was scored to ensure its consistency with the maintenance culture. For four consecutive days, single worms were transferred to fresh plates every 24 hours. Starting from day 5, worms were kept on the same plate for two more days and then killed. This provides a daily readout for the first four days and a day 5 readout representing the last three days combined. This experimental design was performed given that approximately 90% of the worm’s self-progeny is produced within the first three days of adulthood. To obtain fecundity counts, all plates were counted for viable progeny after five days from transferring the mother, thus acquiring both daily and overall self-fecundity. Plates were kept at 20°C across all steps of the experimental design (*SI Appendix*; Fig. S1a).

**Developmental speed measure**

From maintenance cultures, J4 animals were isolated to avoid any outcrossing of the hermaphrodite worms with spontaneous males in the population. After 24 hours, these worms are developed into breeding adults. Afterwards, 10 breeding adults were placed on a fresh plate with 100µl of the respective food source. Plates were incubated for two hours in order to obtain eggs before the mothers were removed. Note that *P. pacificus* lays its eggs in the 2 or 4-cell stage, resulting in at least 40-50 highly synchronized egg clutches. After 75 hours, worms were observed to determine the developmental stage of the progeny. For each strain, and accordingly for each food condition; 40 − 50 mothers were isolated representing 4-5 biological replicates. Between 232 to 489 progenies were staged for each experiment. Plates were kept at 20°C across all steps of the experimental design (*SI Appendix*; Fig. S1b). The time point of 75hrs was chosen to capture the transition rate from juvenile stages to adulthood (Sommer 2015; Sun *et al.* 2021).

**Predation assays: corpse assay**

Two types of predation assays were performed in this study; inter-specific and intra-specific predation assays. In the inter-specific predation assays, young adult *P. pacificus* predators, prey on *C. elegans* L1 larvae; while in the intra-specific predation assays, young adult predators of a particular *P. pacificus* strain prey on *P. pacificus* J2 larvae of the other strain.

Corpse assays were performed to quantify both inter as well as intra-specific predation rates. All assays were conducted as previously described in (Lightfoot *et al.* 2019). In short, for the inter-specific corpse assay, freshly starved *C. elegans* plates were washed with M9 buffer to collect L1 larvae, passed through two Millipore 20µm filters to remove other developmental stages, and followed by centrifugation at 377g/2min to obtain a concentrated larval pellet. One µl of the L1 wash was added onto an empty 6cm (NGM) plate, which represents roughly 3000 larval prey. The *C. elegans* larvae were given at least 1hour window to proportionally spread across the plate. For predators, five *P. pacificus* young adults were blindly picked (independent of mouth-form) from *E. coli* OP50 maintenance cultures. This procedure reflects the predation rates of a population in relevance to mouth-form ratio. Predators were first kept for 10-15 min on an empty plate to reduce body-attached bacteria and were then added to assay plates. The number of corpses was scored after 2 hours with three biological replicates conducted for each assay. For intra-specific predation, we increased the number of predators from 5 to 20 and the assay time from 2 hours to 24 hours as previously reported in (Wilecki *et al.* 2015). In addition, predators were grown on the relevant food source before being blindly picked; i.e., *E. coli* or *Novosphingobium*. For the intra-specific setup, five biological replicates were conducted per assay (*SI Appendix*; Fig. S1c).

#### Details of the model

To model the dynamics of *P. pacificus* in different environments based on our laboratory data, we envisioned the development of a worm as a finite-state Markov chain. The Markov chain is used to construct a stage-structured population model (for more on this approach to modelling population dynamics, see (Nathan Keyfitz 2005; Caswell 2019)). The projection matrix for this chain is:

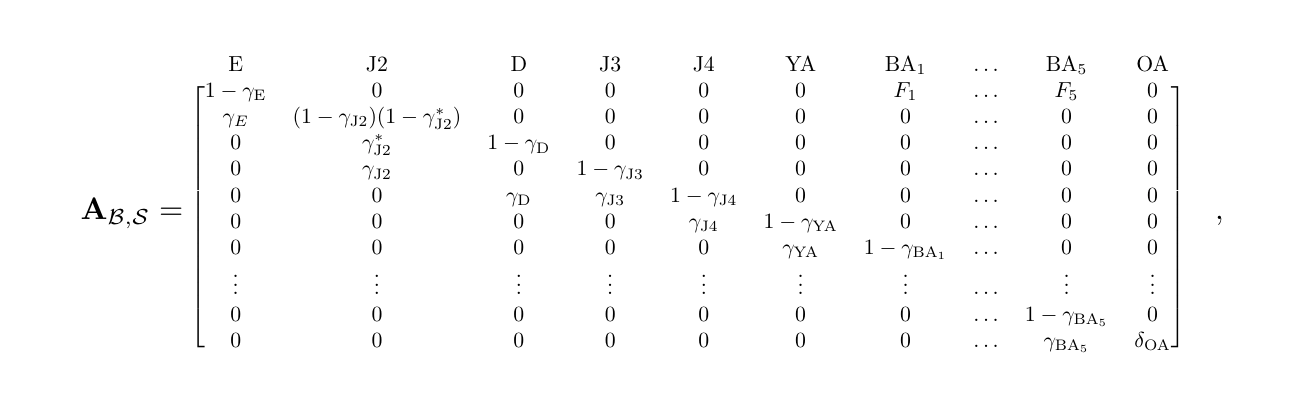

where $\gamma_{i}$ is the probability of transition from developmental stage $i$ into the next developmental stage. In the case of J2, $\gamma_{\text{J2}}$ and $\gamma_{\text{J2}}^{*}$ are the probabilities for J2 → J3 and J2 → D, respectively. Note that as long as food is available ($S_{t}$ > 0), J2 → D transition has a zero probability. We assume that all individuals in each stage survive and develop into the next stage, except for old adults (OA), which have a survival probability, $\delta_{\text{OA}}$. In the absence of food, the transition probabilities of all the juvenile stages, as well as the eggs, are zero, while the transition probability for J2 → D is no longer zero ($\gamma_{\text{J2}}^{*}$ =0.1). The five breeding adult stages (BA_1_ to BA_5_) each have their own respective per capita fecundity, $F_{i}$, for a given bacterial diet (B), based on the daily self-fecundity experiment. For a given *P. pacificus* strain, the transition probabilities and fecundities in the projection matrix depend on the experimentally-informed estimates. The transition probabilities and fecundities for the plastic and the non-plastic are listed in **Table** 1.

| **Transition probabilities** | | | |
| --- | --- | --- | --- |
|  | *E. coli* | *Novosphingobium* | Starvation |
| E > J2 | 0.0415 | 0.0415 | 0 |
| J2 > dauer | 0 | 0 | 0.1 |
| J2 > J3 | 0.055 | 0.055 | 0 |
| J3 > J4 | 0.085 | 0.085 | 0 |
| Dauer > J4 | 0.1 | 0.1 | 0 |
| J4 > YA | 0.07 | 0.07 | 0 |
| YA > BA_i_ | 0.1 | **0.13*, 0.4**** | 0 |
| BA_i_ > BA_i+1_ | 0.0415 | 0.0415 | 0.0415 |
| **Fecundities** | | | |
|  | *E. coli* | *Novosphingobium* | |
| BA_1_ | 22.65*, 19.8** | 11.66*, 16.88** | |
| BA_2_ | 68.45*, 60.3** | 62.53*, 80.77** | |
| BA_3_ | 57.05*, 43.02** | 47.13*, 77.7** | |
| BA_4_ | 33.4*, 19.9** | 13.94*, 16.28** | |
| BA_5_ | 4.97*, 6.6** | 0.72*, 1.4** | |

**Table 1: Parameters used in the model**. The fecundity values are based on the average daily number of eggs laid by a given strain on a given food source. The plastic strain specific values is indicated by * and the non-plastic specific values is indicated by **.

The transition probabilities between different stages are set such that the occupancy time for each of the Markov states in our life cycle, *i.e.,* the average time spent over an individuals' life in that state, would correspond to the developmental speed of *P. pacificus* in hours. The mean occupancy time is obtained by calculating the fundamental matrix (**N**) for transition matrix **U**, where $\mathbf{N}\boldsymbol{=}\left( \mathbf{I-U} \right)^{\mathbf{-1}}$. We used a reduced form of our projection matrix that excluded the dauer stage to calculate the fundamental matrix. On *E. coli*, we assume no difference in developmental speed between RSC017 and RS5405. The first column of the fundamental matrix for these strains on OP50 is [24*.*1*,*18*.*2*,*11*.*8*,*14*.*3*,*10*,*24*,*24*,*24*,*24*,*24*,*200], implying that an egg spends on average 24.1 hours in the egg stage, 18.2 in J2, 11.8 in J3, 14.3 in J4, 10 in YA, 24 in each of the five breeding adult stages, and 200 hours (roughly 8.5 days) in the old adult stage before dying. These values are in line with the laboratory measurements of developmental speed (Sommer 2015; Sun *et al.* 2021). We adjusted the probability of YA → BA_1_ such that the duration of YA stage on *Novosphigobium* would reduce to ≈ 8 and ≈ 6 hours for RSC017 and RS5405, respectively.

**Consumption**

Resource consumption is included in the model by assuming fixed consumption rates for each developmental stage. Given food source S*_t_*, if there exist *m* developmental stages in the population at *t* and *n_i_* individuals belong to developmental stage *i*, the amount of available food in the next step will be:

$$\mathcal{S}_{\mathcal{t+}\boldsymbol{1}}\boldsymbol{=}\mathcal{S}_{\mathcal{t}}\boldsymbol{-}\sum_{\boldsymbol{i=1}}^{\boldsymbol{m}} \boldsymbol{\rho}_{\boldsymbol{i}}\boldsymbol{n}_{\boldsymbol{i}}\quad\boldsymbol{,}$$

(1)

where $\rho_{i}$ is the per capita consumption rate for developmental stage *i*.

**Predation**

If a population consists of two strains, *i* and *j*, the number of surviving J2 individuals of strain *i* at time *t* + 1 is:

$$\boldsymbol{V}_{\boldsymbol{i}}\left( \boldsymbol{t+1} \right)\boldsymbol{=}\boldsymbol{V}_{\boldsymbol{i}}\left( \boldsymbol{t} \right)\boldsymbol{-}\boldsymbol{\eta}_{\boldsymbol{ji}}\boldsymbol{P}_{\boldsymbol{j}}\left( \boldsymbol{t} \right)\boldsymbol{V}_{\boldsymbol{i}}\left( \boldsymbol{t} \right)\quad\boldsymbol{,}$$

(2)

where $\eta_{ji}$ is the rate at which adults from strain $j$ kill J2s of strain $i$, $P_{j}(t)$ is the

number of predatory adults of strain $j$ in the population at time *t*, and $V_{i}(t)$ is the number of J2s of strain $i$. For the plastic strain, the expected number of predatory adults equals the number of adults in the population multiplied by the probability of developing the predatory mouth form on a given bacterial diet.

**Population dynamic**

Assume **n***_i_*(*t*) to be a 12 × 1 array, where each entry represents the count of each developmental stage of strain *i* in a population at time *t*. The expected composition of the population at time *t* + 1 would be:

$$\boldsymbol{n}_{\boldsymbol{i}}\left( \boldsymbol{t+1} \right)\boldsymbol{=}\boldsymbol{A}_{\boldsymbol{t}}\boldsymbol{n}_{\boldsymbol{i}}\left( \boldsymbol{t} \right)\boldsymbol{-}\boldsymbol{K}_{\boldsymbol{i}}\left( \boldsymbol{t} \right)\boldsymbol{-}\boldsymbol{E}_{\boldsymbol{i}}\left( \boldsymbol{t} \right)\quad\boldsymbol{,}$$

(3)

where $K_{i}\left( t \right)$ is the number of J2 individuals of strain $i$ that were killed at time *t* and $E_{i}\left( t \right)$ is the number of dauer larvae that emigrated from the population at time *t*.

**Structured population in one dimension**

In order to investigate the effect of dispersal on the population dynamics, we constructed a one-dimensional structured population that consisted of *n* localities arranged in a line. At each step, a proportion $\omega$of the dauer larvae from a locality emigrates to its neighboring locality if the neighboring locality has more available food, resulting in a one-way dispersal pattern from a source to a sink. Throughout the model, n = 12 and $\omega$= 0*.*1.

**Estimating the predation parameter**

To estimate the predation parameter for $\eta_{ji}$, we fitted the solution to the difference equation 2,

$$\boldsymbol{V}_{\boldsymbol{i}}\left( \boldsymbol{t} \right)\boldsymbol{=}\boldsymbol{V}_{\boldsymbol{i}}\left( \boldsymbol{0} \right)\left( \boldsymbol{1-}\boldsymbol{\eta}_{\boldsymbol{ji}}\boldsymbol{P}_{\boldsymbol{j}}\left( \boldsymbol{t} \right) \right)^{\boldsymbol{t}}$$

(4)

to our empirical data from killing assays. Each killing assay starts with ≈ 3000 J2 worms of strain i ($V_{i}\left( 0 \right)$= 3000) and 20 adults of strain *j*. The number of corpses is counted after 24 hours. Assuming a fixed killing rate over the duration of the killing assay, we estimated the $\eta_{ji}$ that would result in the number of corpses observed in our assay.

#### Details of the statistical analyses

To analyze the experimental data, instead of taking the Frequentist approach, we opted for Bayesian alternatives. To calculate the probability of developing the predatory mouth morph, we assumed the number of observed predatory worms in a sample of n worms follows the likelihood function y$\sim Bernnoulli(\theta)$, where, as our prior, we assume $\theta$ is drawn from a beta distribution with $\alpha=\beta=1$, which corresponds to a uniform distribution. For our Bayesian estimation for comparing two groups, equivalent to t-test, for two samples, a and b, we follow Kruschke’s BEST approach (Kruschke 2013, 2015): we define likelihood functions, $y_{i}^{a}\mathcal{\sim T(}\nu,\mu_{a},\sigma_{a})$ and $y_{i}^{b}\mathcal{\sim T(}\nu,\mu_{b},\sigma_{b})$. As our prior, we assume that the mean of each sample is from a normal distribution, with the mean and twice the standard deviation of a pooled sample. For the standard deviation, we assume a wide uniform prior, $Unifrom(1,300)$. Following Kruschke, we use $\nu=30$; at higher values of $\nu$, the t-distribution converges to the normal distribution. Such an approach is preferable to the standard t-test, since it compares means and standard deviations between to the two groups. The mean and the 95% highest density interval of our estimates of the parameters of interests, difference in means and difference in standard deviations of two groups, as well as the effects size are reported. This approach lacks the simple and, somewhat deceptive, clarity of Null hypothesis significance testing, but the Bayesian approach is more scientifically appealing, and it enables side-stepping the many issues with p-value (Wasserstein & Lazar 2016). All statistical analyses were conducted with PyMC3 in Python 3.10.4, using the No-U-Turn Sampler. In every analysis, we used effective sample size of $\geq10,000$ for stable estimates of HDIs and ensured that all the 4 MCMC chain had converged, i.e., $\hat{R}=1$ (Kruschke 2021). The code used to analyze the data with PyMC3 3.11.4 (Salvatier *et al.* 2016) are included in Jupiter notebooks and are accessible on our Github repository associated with this manuscript. The detailed results of the BEST approach can be found in Tables S6-S8.

4.

Stiernagle, T. Maintenance of *C. elegans*. In: *WormBook* (ed. Community, TCeR).

5.

Sommer, R.J. (2015). *Pristionchus pacificus: A Nematode Model for Comparative and Evolutionary Biology*. Brill.

6.

Sun, S., Rödelsperger, C. & Sommer, R.J. (2021). Single worm transcriptomics identifies a developmental core network of oscillating genes with deep conservation across nematodes. *Genome Res.*, 31, 1590-1601.

7.

Lightfoot, J.W., Wilecki, M., Rödelsperger, C., Moreno, E., Susoy, V., Witte, H. *et al.* (2019). Small peptide-mediated self-recognition prevents cannibalism in predatory nematodes. *Science*, 364, 86-89.

8.

Wilecki, M., Lightfoot, J.W., Susoy, V. & Sommer, R.J. (2015). Predatory feeding behaviour in Pristionchus nematodes is dependent on a phenotypic plasticity and induced by serotonin. *J. Exp. Biol.*, 218, 1306-1313.

9.

Nathan Keyfitz, H.C. (2005). *Applied Mathematical Demography*. Springer New York, NY.

10.

Caswell, H. (2019). *Sensitivity Analysis: Matrix Methods in Demography and Ecology*. Springer Cham.

11.

Kruschke, J.K. (2013). Bayesian estimation supersedes the t test. *J. Exp. Psychol. Gen.*, 142, 573-603.

12.

Kruschke, J.K. (2015). Chapter 16 - Metric-Predicted Variable on One or Two Groups. In: *Doing Bayesian Data Analysis (Second Edition)* (ed. Kruschke, JK). Academic Press Boston, pp. 449-475.

13.

Wasserstein, R.L. & Lazar, N.A. (2016). The ASA Statement on *p*-Values: Context, Process, and Purpose. *Am. Stat.*, 70, 129-133.

14.

Kruschke, J.K. (2021). Bayesian Analysis Reporting Guidelines. *Nat. Hum. Behav.*, 5, 1282-1291.

15.

Salvatier, J., Wiecki, T.V. & Fonnesbeck, C. (2016). Probabilistic programming in Python using PyMC3. *PeerJ Comput. Sci.*, 2, e55.
