## Supplementary figures and images for "Experimental and theoretical support for costs of plasticity and phenotype in a nematode cannibalistic trait"

### Fig S1

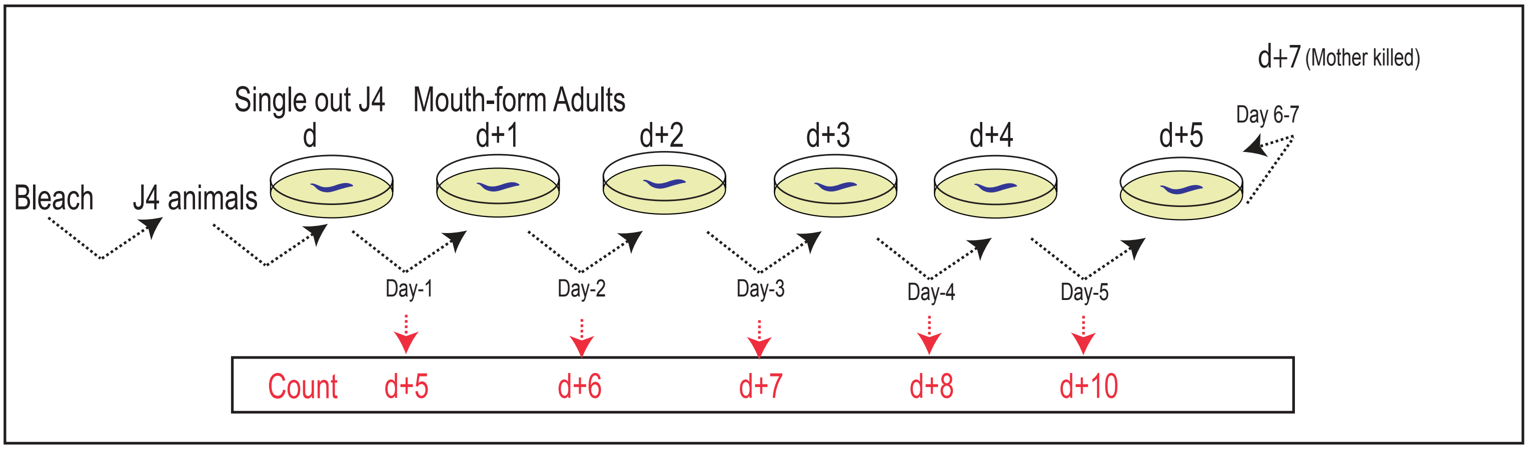

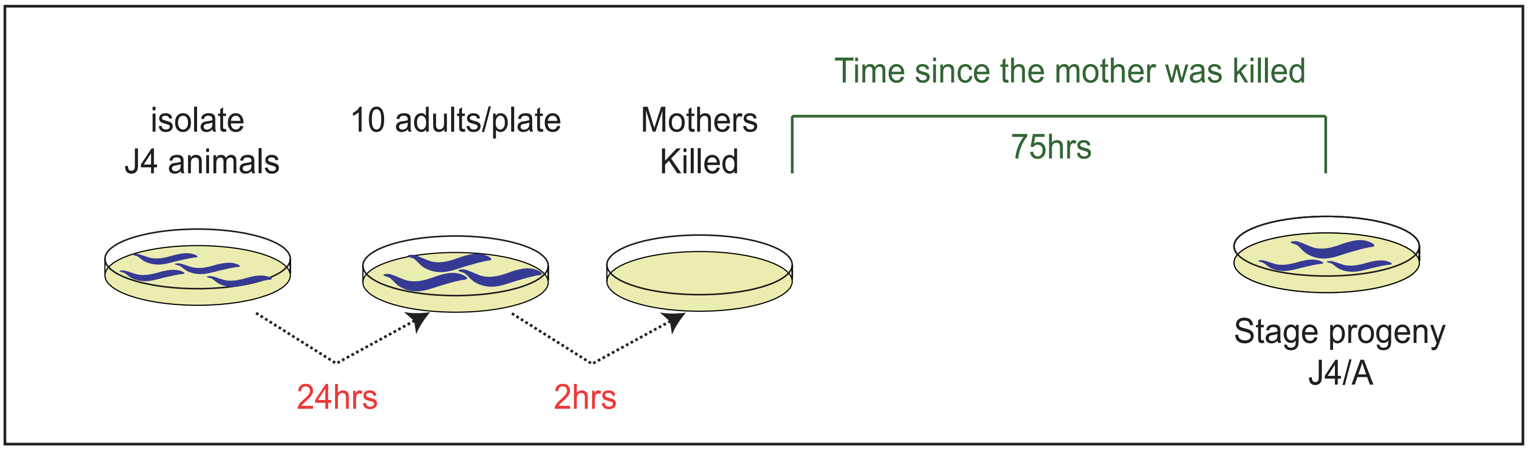

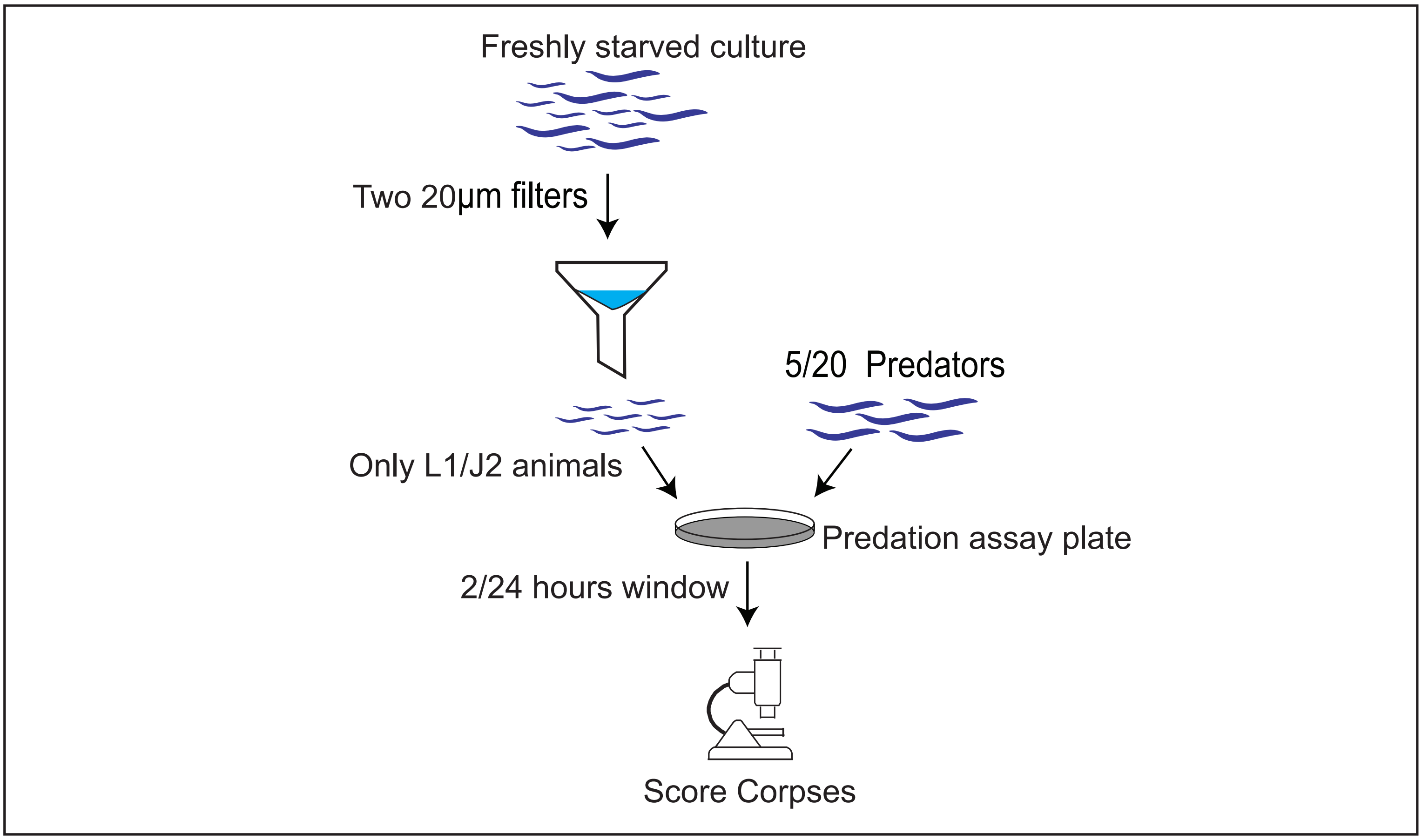

### Fig S2

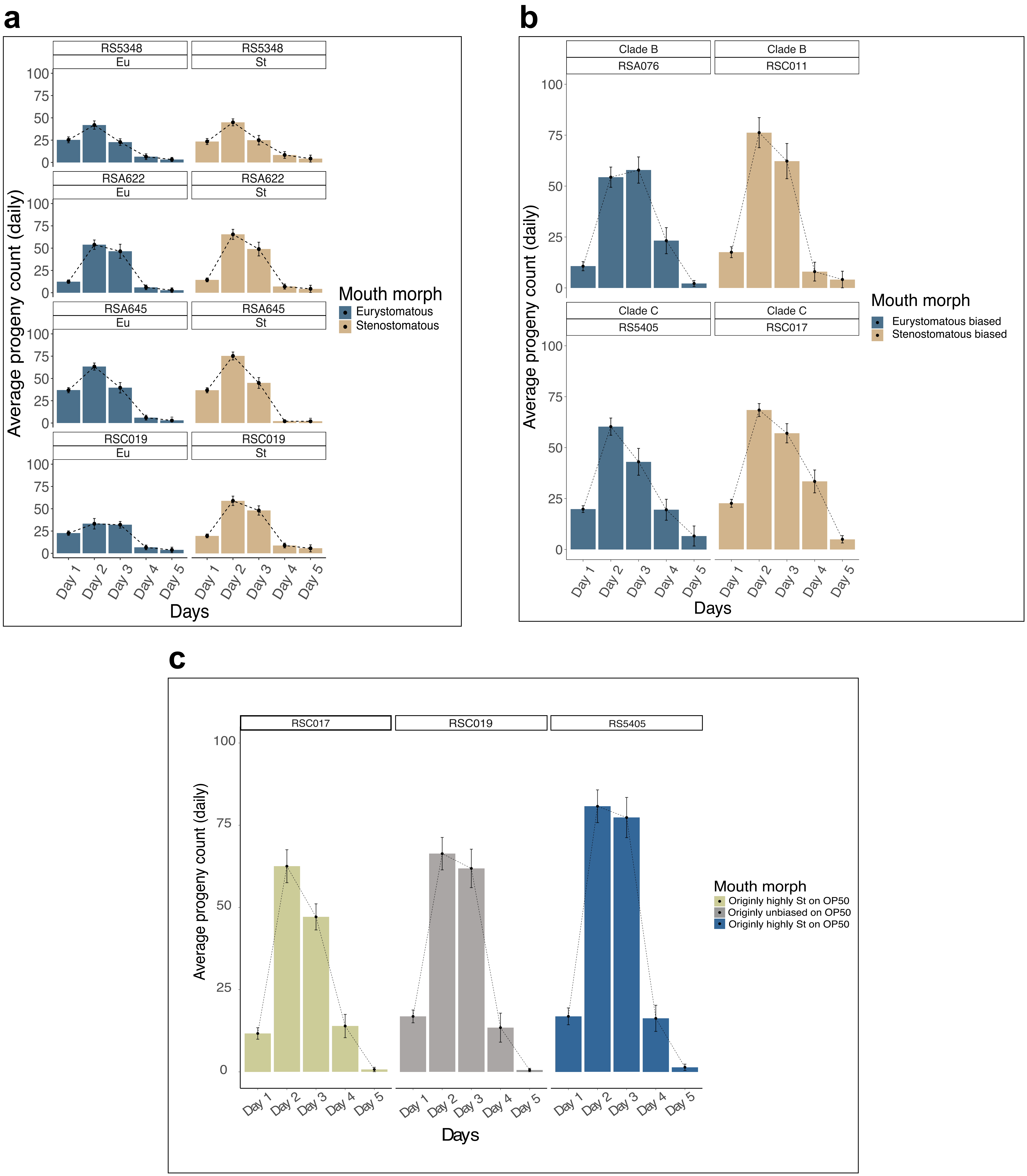

### Fig S3

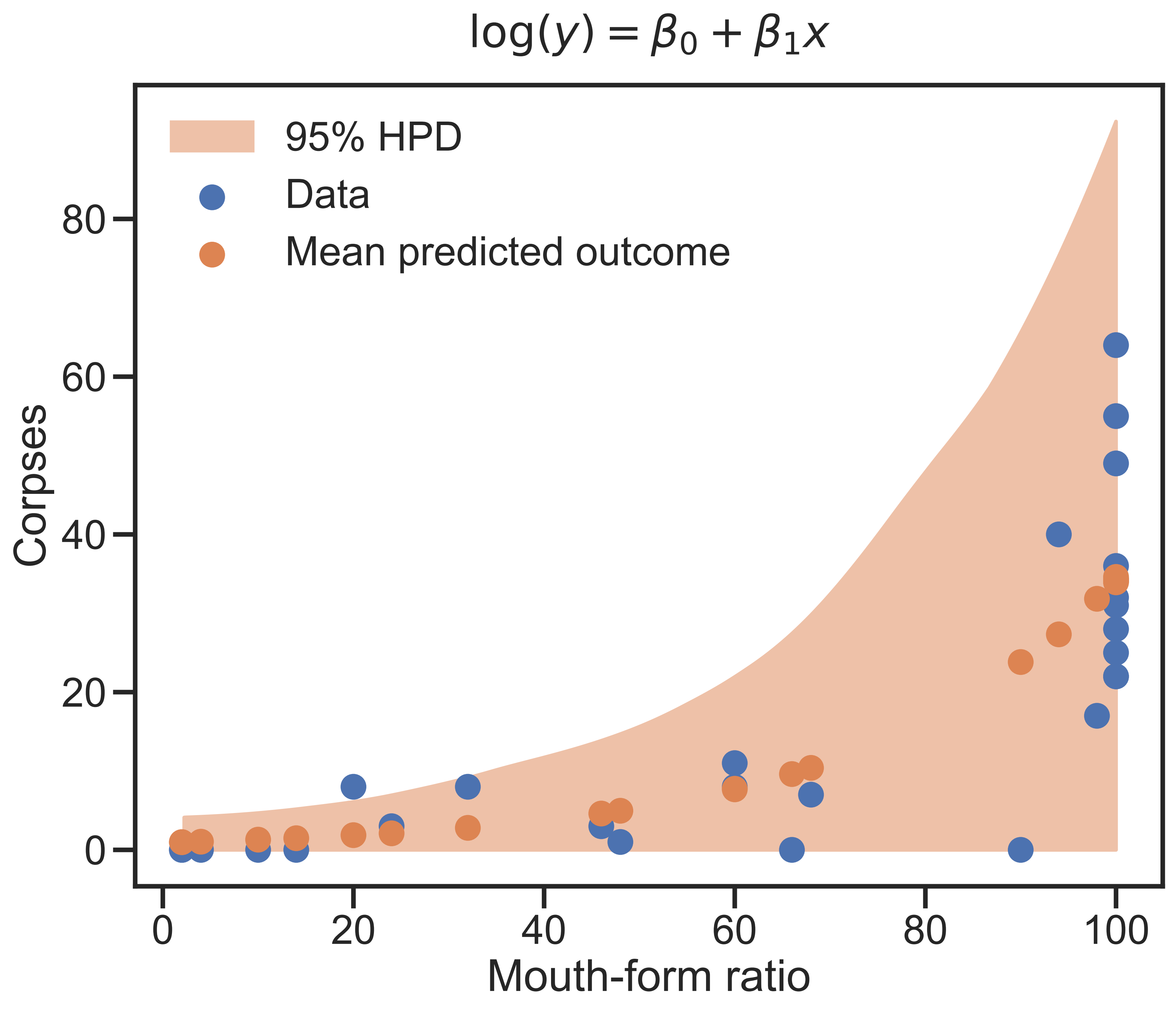

### Fig S4

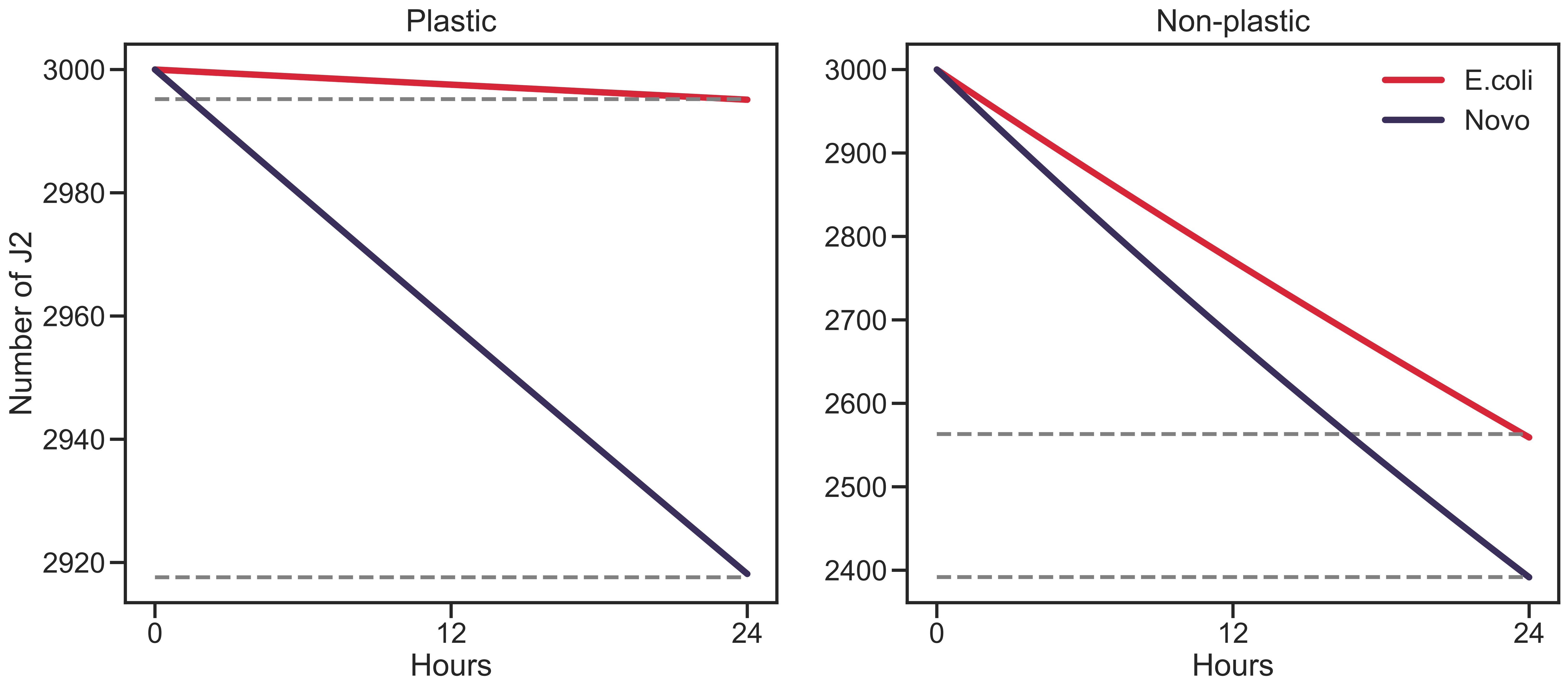

### Fig S5

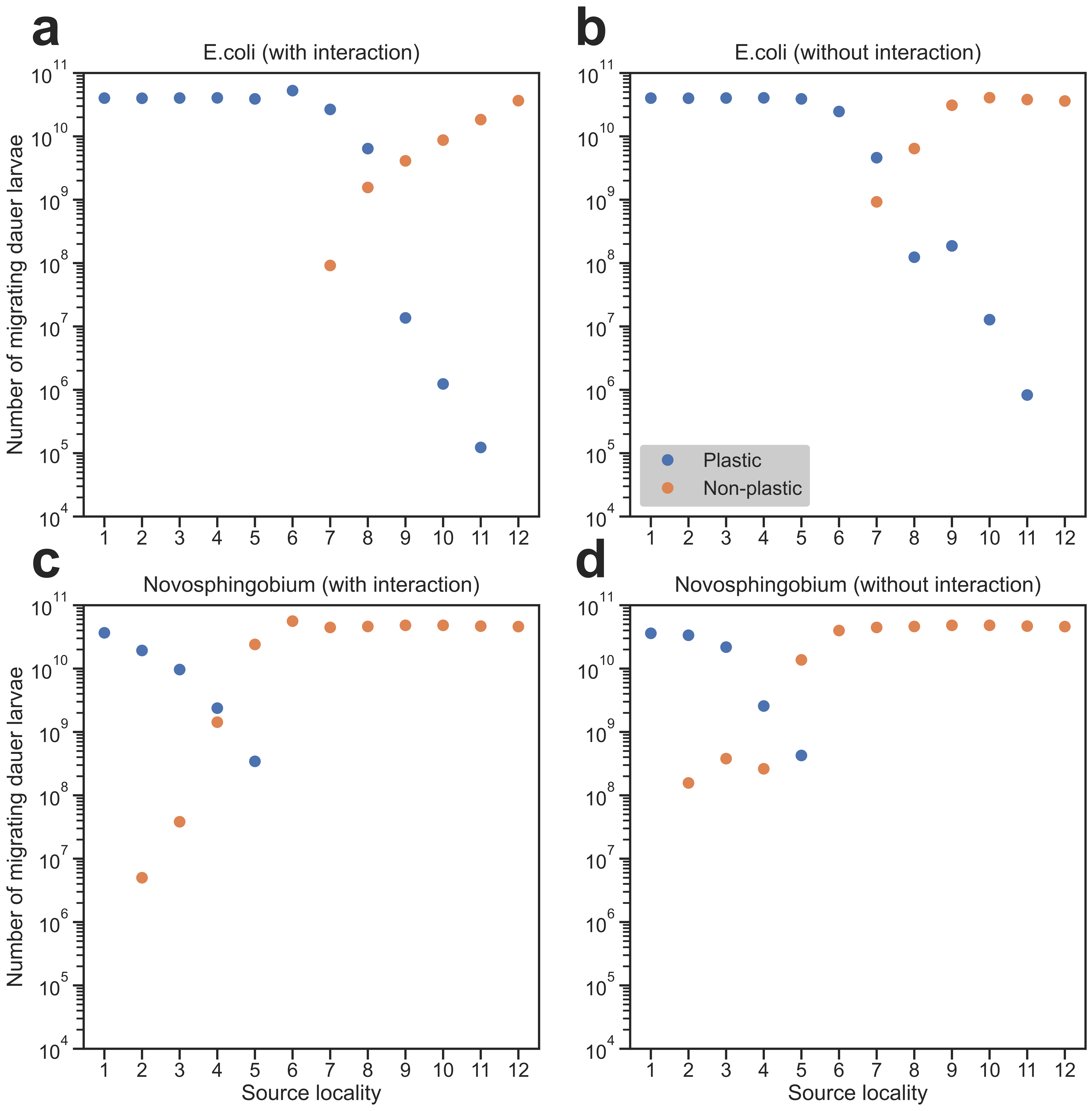

### Fig S6

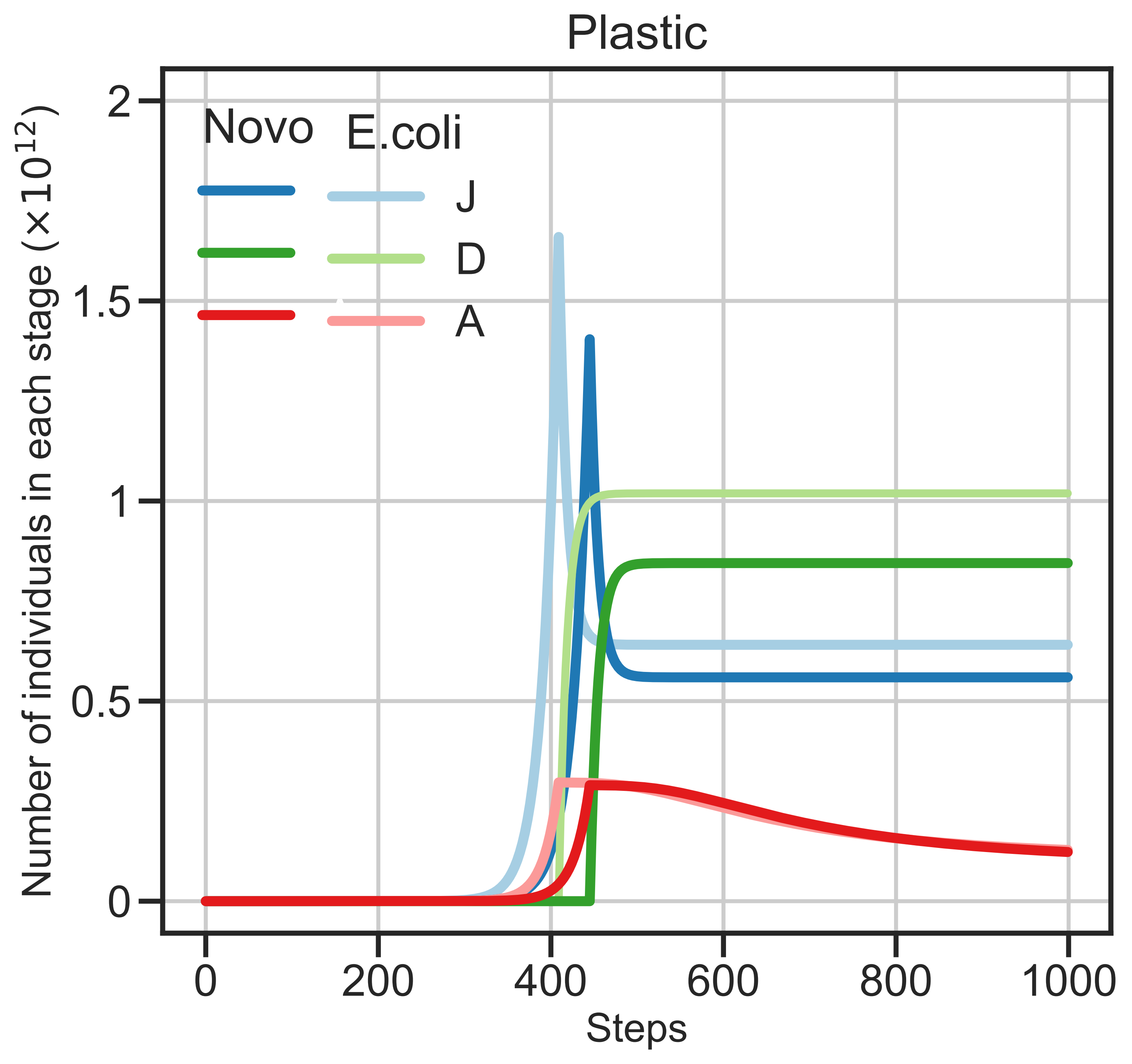
