## Supplementary material for "Experimental and theoretical support for costs of plasticity and phenotype in a nematode cannibalistic trait": Table S1

| Strain | Location/clade | Number of Eu individuals | Number of St individuals | Overall fecundity mean Eu | Overall fecundity mean St | Standard deviation Eu | Standard deviation St | Expirmental setup | Condition |
|---|---|---|---|---|---|---|---|---|---|
| RS5348 | La Reunion, Trois Bassins (TB)/C | 44 | 33 | 99,75 | 105,9393939 | 29,03576383 | 30,79259184 | intra-genotype cost of phenotype | *E. coli* |
| RSA113 | La Reunion, Trois Bassins (TB)/C | 54 | 41 | 68,18518519 | 72,92682927 | 25,92738341 | 29,5375949 | intra-genotype cost of phenotype | *E. coli* |
| RSA622 | Mauritius, Sugarcane Institute (MU)/C | 50 | 49 | 121,5833333 | 140,0169492 | 45,12216625 | 40,81180168 | intra-genotype cost of phenotype | *E. coli* |
| RSA645 | Mauritius, Lakaz Chamarel Med.pla (MU)/A | 55 | 54 | 148,6363636 | 160,8333333 | 38,48056924 | 40,74344489 | intra-genotype cost of phenotype | *E. coli* |
| RSC019 | La Reunion, Colorado (CO)/C | 54 | 53 | 98,2037037 | 140,8490566 | 36,15030448 | 39,93330797 | intra-genotype cost of phenotype | *E. coli* |
| RSC033 | La Reunion, Grand Etang Lake-3 (GE)/C | 54 | 47 | 97,92592593 | 125,5744681 | 36,31302685 | 48,02433472 | intra-genotype cost of phenotype | *E. coli* |
| RSD029 | La Reunion, Nez de Boeuf (NB)/B | 46 | 54 | 143,3478261 | 146,7407407 | 31,13784221 | 32,59365906 | intra-genotype cost of phenotype | *E. coli* |

| Strain | Location/clade | Number of Eu individuals | Number of St individuals | Overall fecundity mean Eu | Overall fecundity mean St | Overall fecundity Eu+St | Standard deviation Eu | Standard deviation St | Standard deviation Eu+St | Expirmental setup | Condition |
|---|---|---|---|---|---|---|---|---|---|---|---|
| RSC011 | La Reunion, Coteau Kerveguen (CK)/ B | 9 | 32 | 173,2222222 | 166,75 | 168,1707317 | 36,04087186 | 45,10131963 | 42,93710659 | inter-genotype cost of phenotype | *E. coli* |
| RSA076 | La Reunion, Nez de Boeuf (NB)/ B | 40 | 0 | 148,3 | 0 | 148,3 | 42,70002702 | 0 | 42,70002702 | inter-genotype cost of phenotype | *E. coli* |
| RS5405 | La Reunion, Trois Bassins (TB)/ C | 40 | 0 | 149,625 | 0 | 149,625 | 45,37348571 | 0 | 45,37348571 | inter-genotype cost of phenotype | *E. coli* |
| RSC017 | La Reunion, Colorado (CO)/ C | 0 | 40 | 0 | 186,525 | 186,525 | 0 | 33,05394775 | 33,05394775 | inter-genotype cost of phenotype | *E. coli* |

| Strain | Location/clade | Number of Eu individuals | Number of St individuals | Overall fecundity mean Eu | Overall fecundity mean St | Overall fecundity Eu+St | Standard deviation Eu | Standard deviation St | Standard deviation Eu+St | Expirmental setup | Condition |
|---|---|---|---|---|---|---|---|---|---|---|---|
| RS54054 | La Reunion, Trois Bassins (TB)/ C | 43 | 0 | 193,0465116 | 0 | 193,0465116 | 33,0273721 | 0 | 33,0273721 | Cost of plasticity | *Novosphingobium* |
| RSC017 | La Reunion, Colorado (CO)/ C | 47 | 0 | 136,2553191 | 0 | 136,2553191 | 27,9887152 | 0 | 27,9887152 | Cost of plasticity | *Novosphingobium* |

### Supplementary Table1
