## Supplementary material for "Experimental and theoretical support for costs of plasticity and phenotype in a nematode cannibalistic trait": Table S2

| Strain | Mean progeny count | Days | Standard deviation | Number of individuals | Condition | Percentage | Animals mouth form | Expirmental setup |
|---|---|---|---|---|---|---|---|---|
| RS5348 | 25,31818182 | Day 1 | 11,19961 | 44 | *E. coli* | 25,3816371 | Eu | intra-genotype cost of phenotype |
| RS5348 | 41,90909091 | Day 2 | 15,71522 | 44 | *E. coli* | 42,0141262 | Eu | intra-genotype cost of phenotype |
| RS5348 | 22,79545455 | Day 3 | 12,53011 | 44 | *E. coli* | 22,852586 | Eu | intra-genotype cost of phenotype |
| RS5348 | 6,431818182 | Day 4 | 10,78414 | 44 | *E. coli* | 6,44793803 | Eu | intra-genotype cost of phenotype |
| RS5348 | 3,29545 | Day 5 | 7,79558 | 44 | *E. coli* | 3,30370927 | Eu | intra-genotype cost of phenotype |
| RS5348 | 23,42424242 | Day 1 | 9,87747 | 33 | *E. coli* | 22,1110036 | St | intra-genotype cost of phenotype |
| RS5348 | 44,9393 | Day 2 | 11,41528 | 33 | *E. coli* | 42,4198574 | St | intra-genotype cost of phenotype |
| RS5348 | 24,93939394 | Day 3 | 15,51801 | 33 | *E. coli* | 23,5412108 | St | intra-genotype cost of phenotype |
| RS5348 | 8,363636364 | Day 4 | 11,01368 | 33 | *E. coli* | 7,89474384 | St | intra-genotype cost of phenotype |
| RS5348 | 4,272727273 | Day 5 | 11,39976 | 33 | *E. coli* | 4,03318435 | St | intra-genotype cost of phenotype |
| RSA645 | 36,83636364 | Day 1 | 11,11349 | 55 | *E. coli* | 24,7828746 | Eu | intra-genotype cost of phenotype |
| RSA645 | 63,34545455 | Day 2 | 14,95081 | 55 | *E. coli* | 42,617737 | Eu | intra-genotype cost of phenotype |
| RSA645 | 39,54545455 | Day 3 | 22,02593 | 55 | *E. coli* | 26,6055046 | Eu | intra-genotype cost of phenotype |
| RSA645 | 6,036363636 | Day 4 | 10,45619 | 55 | *E. coli* | 4,06116208 | Eu | intra-genotype cost of phenotype |
| RSA645 | 2,872727273 | Day 5 | 14,02650 | 55 | *E. coli* | 1,93272171 | Eu | intra-genotype cost of phenotype |
| RSA645 | 36,74074074 | Day 1 | 10,71621 | 54 | *E. coli* | 22,8439839 | St | intra-genotype cost of phenotype |
| RSA645 | 75,22222222 | Day 2 | 16,58104 | 54 | *E. coli* | 46,7702936 | St | intra-genotype cost of phenotype |
| RSA645 | 44,92592593 | Day 3 | 22,62979 | 54 | *E. coli* | 27,9332182 | St | intra-genotype cost of phenotype |
| RSA645 | 2,018518519 | Day 4 | 5,06371 | 54 | *E. coli* | 1,25503742 | St | intra-genotype cost of phenotype |
| RSA645 | 1,925925926 | Day 5 | 12,64143 | 54 | *E. coli* | 1,1974669 | St | intra-genotype cost of phenotype |
| RSA622 | 12,4 | Day 1 | 6,99927 | 50 | *E. coli* | 10,1987663 | Eu | intra-genotype cost of phenotype |
| RSA622 | 53,86666667 | Day 2 | 18,98391 | 50 | *E. coli* | 44,304318 | Eu | intra-genotype cost of phenotype |
| RSA622 | 46,46666667 | Day 3 | 28,50600 | 50 | *E. coli* | 38,2179575 | Eu | intra-genotype cost of phenotype |
| RSA622 | 6 | Day 4 | 9,42949 | 50 | *E. coli* | 4,93488691 | Eu | intra-genotype cost of phenotype |
| RSA622 | 2,85 | Day 5 | 9,37329 | 50 | *E. coli* | 2,34407128 | Eu | intra-genotype cost of phenotype |
| RSA622 | 14,37288136 | Day 1 | 7,74351 | 49 | *E. coli* | 10,2651011 | St | intra-genotype cost of phenotype |
| RSA622 | 65,42372881 | Day 2 | 20,17271 | 49 | *E. coli* | 46,725578 | St | intra-genotype cost of phenotype |
| RSA622 | 48,96610169 | Day 3 | 27,42071 | 49 | *E. coli* | 34,9715531 | St | intra-genotype cost of phenotype |
| RSA622 | 6,898305085 | Day 4 | 10,12626 | 49 | *E. coli* | 4,92676431 | St | intra-genotype cost of phenotype |
| RSA622 | 4,355932203 | Day 5 | 14,28860 | 49 | *E. coli* | 3,11100351 | St | intra-genotype cost of phenotype |
| RSC019 | 22,61111111 | Day 1 | 9,51546 | 54 | *E. coli* | 23,024703 | Eu | intra-genotype cost of phenotype |
| RSC019 | 33,18518519 | Day 2 | 22,06515 | 54 | *E. coli* | 33,7921931 | Eu | intra-genotype cost of phenotype |
| RSC019 | 32,01851852 | Day 3 | 13,87340 | 54 | *E. coli* | 32,6041863 | Eu | intra-genotype cost of phenotype |
| RSC019 | 6,611111111 | Day 4 | 9,97434 | 54 | *E. coli* | 6,73203847 | Eu | intra-genotype cost of phenotype |
| RSC019 | 3,777777778 | Day 5 | 11,58344 | 54 | *E. coli* | 3,84687913 | Eu | intra-genotype cost of phenotype |
| RSC019 | 19,52830189 | Day 1 | 7,97015 | 53 | *E. coli* | 13,8647019 | St | intra-genotype cost of phenotype |
| RSC019 | 58,88679245 | Day 2 | 19,53071 | 53 | *E. coli* | 41,8084394 | St | intra-genotype cost of phenotype |
| RSC019 | 48,05660377 | Day 3 | 19,27475 | 53 | *E. coli* | 34,119223 | St | intra-genotype cost of phenotype |
| RSC019 | 8,773584906 | Day 4 | 9,97434 | 53 | *E. coli* | 6,22906899 | St | intra-genotype cost of phenotype |
| RSC019 | 5,603773585 | Day 5 | 14,41999 | 53 | *E. coli* | 3,97856664 | St | intra-genotype cost of phenotype |
| RSC011 | 17,53658537 | Day 1 | 8,553062495 | 41 | *E. coli* | 10,4278463 | highly st | inter-genotype cost of phenotype |
| RSC011 | 76,24390244 | Day 2 | 23,9629923 | 41 | *E. coli* | 45,3372009 | highly st | inter-genotype cost of phenotype |
| RSC011 | 62,26829268 | Day 3 | 27,51064081 | 41 | *E. coli* | 37,026831 | highly st | inter-genotype cost of phenotype |
| RSC011 | 8 | Day 4 | 14,91978552 | 41 | *E. coli* | 4,75707034 | highly st | inter-genotype cost of phenotype |
| RSC011 | 4,12195122 | Day 5 | 13,14000594 | 41 | *E. coli* | 2,45105149 | highly st | inter-genotype cost of phenotype |
| RSA076 | 10,675 | Day 1 | 7,230233886 | 40 | *E. coli* | 7,1982468 | highly Eu | inter-genotype cost of phenotype |
| RSA076 | 54,375 | Day 2 | 16,02031963 | 40 | *E. coli* | 36,6655428 | highly Eu | inter-genotype cost of phenotype |
| RSA076 | 57,875 | Day 3 | 21,06624728 | 40 | *E. coli* | 39,0256237 | highly Eu | inter-genotype cost of phenotype |
| RSA076 | 23,2 | Day 4 | 20,65491808 | 40 | *E. coli* | 15,6439649 | highly Eu | inter-genotype cost of phenotype |
| RSA076 | 2,175 | Day 5 | 4,684330125 | 40 | *E. coli* | 1,46662171 | highly Eu | inter-genotype cost of phenotype |
| RSC017 | 22,65 | Day 1 | 6,290204024 | 40 | *E. coli* | 12,1431444 | highly st | inter-genotype cost of phenotype |
| RSC017 | 68,45 | Day 2 | 10,19684797 | 40 | *E. coli* | 36,6974936 | highly st | inter-genotype cost of phenotype |
| RSC017 | 57,05 | Day 3 | 15,19606899 | 40 | *E. coli* | 30,5857124 | highly st | inter-genotype cost of phenotype |
| RSC017 | 33,4 | Day 4 | 18,09348941 | 40 | *E. coli* | 17,9064469 | highly st | inter-genotype cost of phenotype |
| RSC017 | 4,975 | Day 5 | 5,757882 | 40 | *E. coli* | 2,66720279 | highly st | inter-genotype cost of phenotype |
| RS5405 | 19,8 | Day 1 | 5,302152441 | 40 | *E. coli* | 13,2696386 | highly Eu | inter-genotype cost of phenotype |
| RS5405 | 60,3 | Day 2 | 13,61597855 | 40 | *E. coli* | 40,4120813 | highly Eu | inter-genotype cost of phenotype |
| RS5405 | 43,025 | Day 3 | 18,94457165 | 40 | *E. coli* | 28,8346567 | highly Eu | inter-genotype cost of phenotype |
| RS5405 | 19,4878049 | Day 4 | 16,20667795 | 40 | *E. coli* | 13,0604105 | highly Eu | inter-genotype cost of phenotype |
| RS5405 | 6,6 | Day 5 | 4,684330125 | 40 | *E. coli* | 4,42321288 | highly Eu | inter-genotype cost of phenotype |
| RSC017 | 11,67391304 | Day 1 | 6,09208721 | 47 | *Novosphingobium* | 8,58419751 | on *E. coli* highly St | cost of plasticity |
| RSC017 | 62,53191489 | Day 2 | 17,5261929 | 47 | *Novosphingobium* | 45,9816949 | on *E. coli* highly St | cost of plasticity |
| RSC017 | 47,12765957 | Day 3 | 13,86835155 | 47 | *Novosphingobium* | 34,6544588 | on *E. coli* highly St | cost of plasticity |
| RSC017 | 13,93617021 | Day 4 | 12,39460657 | 47 | *Novosphingobium* | 10,2477068 | on *E. coli* highly St | cost of plasticity |
| RSC017 | 0,723404255 | Day 5 | 2,337800226 | 47 | *Novosphingobium* | 0,53194203 | on *E. coli* highly St | cost of plasticity |
| RS5405 | 16,88372093 | Day 1 | 8,555725742 | 43 | *Novosphingobium* | 8,76315975 | on *E. coli* highly Eu | cost of plasticity |
| RS5405 | 80,76744186 | Day 2 | 16,65595448 | 43 | *Novosphingobium* | 41,9207353 | on *E. coli* highly Eu | cost of plasticity |
| RS5405 | 77,34146341 | Day 3 | 20,45382126 | 43 | *Novosphingobium* | 40,1425493 | on *E. coli* highly Eu | cost of plasticity |
| RS5405 | 16,27906977 | Day 4 | 13,30471735 | 43 | *Novosphingobium* | 8,44932759 | on *E. coli* highly Eu | cost of plasticity |
| RS5405 | 1,395348837 | Day 5 | 3,193293082 | 43 | *Novosphingobium* | 0,72422808 | on *E. coli* highly Eu | cost of plasticity |

**Supplementary Table2**
