## Supplementary material for "Experimental and theoretical support for costs of plasticity and phenotype in a nematode cannibalistic trait": Table S3

| Strain | Total count | J1(eggs) | J2 | J3 | J4 | Young Adult (YA) | Adult with eggs(BA) | Number of Mothers | %J1(eggs) | %J2 | %J3 | % J4 | %YA | %BA | Condition |
|---|---|---|---|---|---|---|---|---|---|---|---|---|---|---|---|
| RSC017 | 70 | 0 | 1 | 0 | 8 | 61 | 0 | 9 | 0 | 1,428571 | 0 | 11,42857 | 87,14286 | 0 | *E. coli* |
| RSC017 | 72 | 0 | 0 | 3 | 6 | 63 | 0 | 9 | 0 | 0 | 4,166667 | 8,333333 | 87,5 | 0 | *E. coli* |
| RSC017 | 66 | 1 | 1 | 1 | 8 | 55 | 0 | 10 | 1,515152 | 1,515152 | 1,515152 | 12,12121 | 83,33333 | 0 | *E. coli* |
| RSC017 | 62 | 0 | 0 | 1 | 0 | 61 | 0 | 9 | 0 | 0 | 1,612903 | 0 | 98,3871 | 0 | *E. coli* |
| RSC017 | 50 | 0 | 0 | 0 | 3 | 47 | 0 | 10 | 0 | 0 | 0 | 6 | 94 | 0 | *E. coli* |
| RS5405 | 69 | 0 | 0 | 0 | 13 | 56 | 0 | 10 | 0 | 0 | 0 | 18,84058 | 81,15942 | 0 | *E. coli* |
| RS5405 | 51 | 0 | 0 | 0 | 10 | 41 | 0 | 10 | 0 | 0 | 0 | 19,60784 | 80,39216 | 0 | *E. coli* |
| RS5405 | 71 | 0 | 1 | 0 | 12 | 58 | 0 | 10 | 0 | 1,408451 | 0 | 16,90141 | 81,69014 | 0 | *E. coli* |
| RS5405 | 56 | 0 | 1 | 0 | 7 | 48 | 0 | 10 | 0 | 1,785714 | 0 | 12,5 | 85,71429 | 0 | *E. coli* |
| RS5405 | 59 | 0 | 0 | 1 | 6 | 52 | 0 | 10 | 0 | 0 | 1,694915 | 10,16949 | 88,13559 | 0 | *E. coli* |
| RSC011 | 68 | 0 | 0 | 1 | 31 | 36 | 0 | 10 | 0 | 0 | 1,470588 | 45,58824 | 52,94118 | 0 | *E. coli* |
| RSC011 | 98 | 0 | 1 | 1 | 33 | 63 | 0 | 9 | 0 | 1,020408 | 1,020408 | 33,67347 | 64,28571 | 0 | *E. coli* |
| RSC011 | 41 | 0 | 0 | 0 | 21 | 20 | 0 | 10 | 0 | 0 | 0 | 51,21951 | 48,78049 | 0 | *E. coli* |
| RSC011 | 56 | 0 | 0 | 0 | 17 | 39 | 0 | 9 | 0 | 0 | 0 | 30,35714 | 69,64286 | 0 | *E. coli* |
| RSA076 | 50 | 0 | 1 | 1 | 38 | 10 | 0 | 10 | 0 | 2 | 2 | 76 | 20 | 0 | *E. coli* |
| RSA076 | 59 | 0 | 1 | 4 | 44 | 10 | 0 | 9 | 0 | 1,694915 | 6,779661 | 74,57627 | 16,94915 | 0 | *E. coli* |
| RSA076 | 70 | 0 | 0 | 1 | 53 | 16 | 0 | 9 | 0 | 0 | 1,428571 | 75,71429 | 22,85714 | 0 | *E. coli* |
| RSA076 | 53 | 0 | 0 | 0 | 26 | 27 | 0 | 9 | 0 | 0 | 0 | 49,0566 | 50,9434 | 0 | *E. coli* |
| RSC017 | 72 | 0 | 0 | 0 | 5 | 55 | 12 | 10 | 0 | 0 | 0 | 6,944444 | 76,38889 | 16,66667 | *Novosphingobium* |
| RSC017 | 87 | 0 | 0 | 0 | 2 | 67 | 18 | 10 | 0 | 0 | 0 | 2,298851 | 77,01149 | 20,68966 | *Novosphingobium* |
| RSC017 | 65 | 0 | 0 | 0 | 6 | 40 | 19 | 9 | 0 | 0 | 0 | 9,230769 | 61,53846 | 29,23077 | *Novosphingobium* |
| RSC017 | 77 | 0 | 0 | 2 | 3 | 47 | 25 | 10 | 0 | 0 | 2,597403 | 3,896104 | 61,03896 | 32,46753 | *Novosphingobium* |
| RS5405 | 124 | 0 | 0 | 0 | 3 | 46 | 75 | 10 | 0 | 0 | 0 | 2,419355 | 37,09677 | 60,48387 | *Novosphingobium* |
| RS5405 | 131 | 0 | 0 | 1 | 1 | 21 | 108 | 10 | 0 | 0 | 0,763359 | 0,763359 | 16,03053 | 82,44275 | *Novosphingobium* |
| RS5405 | 126 | 0 | 0 | 0 | 0 | 13 | 113 | 10 | 0 | 0 | 0 | 0 | 10,31746 | 89,68254 | *Novosphingobium* |
| RS5405 | 108 | 0 | 0 | 0 | 4 | 12 | 92 | 10 | 0 | 0 | 0 | 3,703704 | 11,11111 | 85,18519 | *Novosphingobium* |

**Supplementary Table3**
