## Supplementary material for "Experimental and theoretical support for costs of plasticity and phenotype in a nematode cannibalistic trait": Table S4

| Strain | Condition | Number of Eu animals | Total number of animals counted | %Eu | Expirmental setup |
|---|---|---|---|---|---|
| RSA133 | *E. coli* | 28 | 47 | 59,57447 | intra-genotype cost of phenotype |
| RSA133 | *E. coli* | 63 | 93 | 67,74194 | intra-genotype cost of phenotype |
| RSA133 | *E. coli* | 67 | 103 | 65,04854 | intra-genotype cost of phenotype |
| RSD029 | *E. coli* | 33 | 47 | 70,21277 | intra-genotype cost of phenotype |
| RSD029 | *E. coli* | 38 | 71 | 53,52113 | intra-genotype cost of phenotype |
| RSD029 | *E. coli* | 40 | 70 | 57,14286 | intra-genotype cost of phenotype |
| RSC033 | *E. coli* | 32 | 45 | 71,11111 | intra-genotype cost of phenotype |
| RSC033 | *E. coli* | 47 | 58 | 81,03448 | intra-genotype cost of phenotype |
| RSC033 | *E. coli* | 40 | 54 | 74,07407 | intra-genotype cost of phenotype |
| RSC019 | *E. coli* | 33 | 50 | 66 | intra-genotype cost of phenotype |
| RSC019 | *E. coli* | 30 | 50 | 60 | intra-genotype cost of phenotype |
| RSC019 | *E. coli* | 23 | 50 | 46 | intra-genotype cost of phenotype |
| RS5348 | *E. coli* | 30 | 53 | 56,60377 | intra-genotype cost of phenotype |
| RS5348 | *E. coli* | 20 | 35 | 57,14286 | intra-genotype cost of phenotype |
| RS5348 | *E. coli* | 30 | 50 | 60 | intra-genotype cost of phenotype |
| RSA645 | *E. coli* | 57 | 111 | 51,35135 | intra-genotype cost of phenotype |
| RSA645 | *E. coli* | 34 | 50 | 68 | intra-genotype cost of phenotype |
| RSA645 | *E. coli* | 26 | 50 | 52 | intra-genotype cost of phenotype |
| RSA622 | *E. coli* | 32 | 116 | 27,58621 | intra-genotype cost of phenotype |
| RSA622 | *E. coli* | 96 | 143 | 67,13287 | intra-genotype cost of phenotype |
| RSA622 | *E. coli* | 16 | 30 | 53,33333 | intra-genotype cost of phenotype |
| RSC017 | *E. coli* | 1 | 50 | 2 | inter-genotype cost of phenotype |
| RSC017 | *E. coli* | 5 | 50 | 10 | inter-genotype cost of phenotype |
| RSC017 | *E. coli* | 1 | 50 | 2 | inter-genotype cost of phenotype |
| RS5405 | *E. coli* | 50 | 50 | 100 | inter-genotype cost of phenotype |
| RS5405 | *E. coli* | 50 | 50 | 100 | inter-genotype cost of phenotype |
| RS5405 | *E. coli* | 50 | 50 | 100 | inter-genotype cost of phenotype |
| RSC011 | *E. coli* | 10 | 50 | 20 | inter-genotype cost of phenotype |
| RSC011 | *E. coli* | 16 | 50 | 32 | inter-genotype cost of phenotype |
| RSC011 | *E. coli* | 24 | 50 | 48 | inter-genotype cost of phenotype |
| RSA076 | *E. coli* | 50 | 50 | 100 | inter-genotype cost of phenotype |
| RSA076 | *E. coli* | 50 | 50 | 100 | inter-genotype cost of phenotype |
| RSA076 | *E. coli* | 50 | 50 | 100 | inter-genotype cost of phenotype |
| RSA619 | *E. coli* | 49 | 50 | 98 | Predation assay |
| RSA619 | *E. coli* | 50 | 50 | 100 | Predation assay |
| RSA619 | *E. coli* | 47 | 50 | 94 | Predation assay |
| RSA639 | *E. coli* | 50 | 50 | 100 | Predation assay |
| RSA639 | *E. coli* | 50 | 50 | 100 | Predation assay |
| RSA639 | *E. coli* | 50 | 50 | 100 | Predation assay |
| RSA635 | *E. coli* | 45 | 50 | 90 | Predation assay |
| RSA635 | *E. coli* | 34 | 50 | 68 | Predation assay |
| RSA635 | *E. coli* | 30 | 50 | 60 | Predation assay |
| RS5200 | *E. coli* | 7 | 50 | 14 | Predation assay |
| RS5200 | *E. coli* | 12 | 50 | 24 | Predation assay |
| RS5200 | *E. coli* | 2 | 50 | 4 | Predation assay |
| RSC017 | *Novosphingobium* | 44 | 50 | 88 | cost of plasticity |
| RSC017 | *Novosphingobium* | 47 | 50 | 94 | cost of plasticity |
| RSC017 | *Novosphingobium* | 42 | 50 | 84 | cost of plasticity |
| RS5405 | *Novosphingobium* | 50 | 50 | 100 | cost of plasticity |
| RS5405 | *Novosphingobium* | 50 | 50 | 100 | cost of plasticity |
| RS5405 | *Novosphingobium* | 50 | 50 | 100 | cost of plasticity |

**Supplementary Table4**
