## Supplementary material for "Experimental and theoretical support for costs of plasticity and phenotype in a nematode cannibalistic trait": Table S5

| Prey | Predator | Corpses | Condition | Experimental setup |
|---|---|---|---|---|
| *C. elegans* (N2) | *P. Pacificus* (RSA076) | 28 | *E. coli* | inter-specific assay |
| *C. elegans* (N2) | *P. Pacificus* (RSA076) | 31 | *E. coli* | inter-specific assay |
| *C. elegans* (N2) | *P. Pacificus* (RSA076) | 49 | *E. coli* | inter-specific assay |
| *C. elegans* (N2) | *P. Pacificus* (RSC011) | 8 | *E. coli* | inter-specific assay |
| *C. elegans* (N2) | *P. Pacificus* (RSC011) | 8 | *E. coli* | inter-specific assay |
| *C. elegans* (N2) | *P. Pacificus* (RSC011) | 1 | *E. coli* | inter-specific assay |
| *C. elegans* (N2) | *P. Pacificus* (RS5405) | 32 | *E. coli* | inter-specific assay |
| *C. elegans* (N2) | *P. Pacificus* (RS5405) | 36 | *E. coli* | inter-specific assay |
| *C. elegans* (N2) | *P. Pacificus* (RS5405) | 25 | *E. coli* | inter-specific assay |
| *C. elegans* (N2) | *P. Pacificus* (RSC017) | 0 | *E. coli* | inter-specific assay |
| *C. elegans* (N2) | *P. Pacificus* (RSC017) | 0 | *E. coli* | inter-specific assay |
| *C. elegans* (N2) | *P. Pacificus* (RSC017) | 0 | *E. coli* | inter-specific assay |
| *C. elegans* (N2) | *P. Pacificus* (RSC019) | 0 | *E. coli* | inter-specific assay |
| *C. elegans* (N2) | *P. Pacificus* (RSC019) | 11 | *E. coli* | inter-specific assay |
| *C. elegans* (N2) | *P. Pacificus* (RSC019) | 3 | *E. coli* | inter-specific assay |
| *C. elegans* (N2) | *P. Pacificus* (RSA639) | 55 | *E. coli* | inter-specific assay |
| *C. elegans* (N2) | *P. Pacificus* (RSA639) | 64 | *E. coli* | inter-specific assay |
| *C. elegans* (N2) | *P. Pacificus* (RSA639) | 22 | *E. coli* | inter-specific assay |
| *C. elegans* (N2) | *P. Pacificus* (RS5200) | 0 | *E. coli* | inter-specific assay |
| *C. elegans* (N2) | *P. Pacificus* (RS5200) | 3 | *E. coli* | inter-specific assay |
| *C. elegans* (N2) | *P. Pacificus* (RS5200) | 0 | *E. coli* | inter-specific assay |
| *C. elegans* (N2) | *P. Pacificus* (RSA635) | 0 | *E. coli* | inter-specific assay |
| *C. elegans* (N2) | *P. Pacificus* (RSA635) | 7 | *E. coli* | inter-specific assay |
| *C. elegans* (N2) | *P. Pacificus* (RSA635) | 8 | *E. coli* | inter-specific assay |
| *C. elegans* (N2) | *P. Pacificus* (RSA619) | 17 | *E. coli* | inter-specific assay |
| *C. elegans* (N2) | *P. Pacificus* (RSA619) | 22 | *E. coli* | inter-specific assay |
| *C. elegans* (N2) | *P. Pacificus* (RSA619) | 40 | *E. coli* | inter-specific assay |
| RSC017 | RSC017 | 0 | *E. coli* | intra-specific assay |
| RSC017 | RSC017 | 0 | *E. coli* | intra-specific assay |
| RSC017 | RSC017 | 0 | *E. coli* | intra-specific assay |
| RSC017 | RSC017 | 0 | *E. coli* | intra-specific assay |
| RSC017 | RSC017 | 0 | *E. coli* | intra-specific assay |
| RSC017 | RS5405 | 443 | *E. coli* | intra-specific assay |
| RSC017 | RS5405 | 539 | *E. coli* | intra-specific assay |
| RSC017 | RS5405 | 340 | *E. coli* | intra-specific assay |
| RSC017 | RS5405 | 496 | *E. coli* | intra-specific assay |
| RSC017 | RS5405 | 366 | *E. coli* | intra-specific assay |
| RS5405 | RSC017 | 7 | *E. coli* | intra-specific assay |
| RS5405 | RSC017 | 4 | *E. coli* | intra-specific assay |
| RS5405 | RSC017 | 10 | *E. coli* | intra-specific assay |
| RS5405 | RSC017 | 0 | *E. coli* | intra-specific assay |
| RS5405 | RSC017 | 3 | *E. coli* | intra-specific assay |
| RS5405 | RS5405 | 0 | *E. coli* | intra-specific assay |
| RS5405 | RS5405 | 0 | *E. coli* | intra-specific assay |
| RS5405 | RS5405 | 0 | *E. coli* | intra-specific assay |
| RS5405 | RS5405 | 0 | *E. coli* | intra-specific assay |
| RS5405 | RS5405 | 1 | *E. coli* | intra-specific assay |
| RSC017 | RSC017 | 0 | *Novosphingobium* | intra-specific assay |
| RSC017 | RSC017 | 0 | *Novosphingobium* | intra-specific assay |
| RSC017 | RSC017 | 0 | *Novosphingobium* | intra-specific assay |
| RSC017 | RSC017 | 0 | *Novosphingobium* | intra-specific assay |
| RSC017 | RSC017 | 0 | *Novosphingobium* | intra-specific assay |
| RSC017 | RS5405 | 587 | *Novosphingobium* | intra-specific assay |
| RSC017 | RS5405 | 691 | *Novosphingobium* | intra-specific assay |
| RSC017 | RS5405 | 720 | *Novosphingobium* | intra-specific assay |
| RSC017 | RS5405 | 530 | *Novosphingobium* | intra-specific assay |
| RSC017 | RS5405 | 513 | *Novosphingobium* | intra-specific assay |
| RS5405 | RSC017 | 102 | *Novosphingobium* | intra-specific assay |
| RS5405 | RSC017 | 90 | *Novosphingobium* | intra-specific assay |
| RS5405 | RSC017 | 42 | *Novosphingobium* | intra-specific assay |
| RS5405 | RSC017 | 87 | *Novosphingobium* | intra-specific assay |
| RS5405 | RSC017 | 91 | *Novosphingobium* | intra-specific assay |
| RS5405 | RS5405 | 0 | *Novosphingobium* | intra-specific assay |
| RS5405 | RS5405 | 0 | *Novosphingobium* | intra-specific assay |
| RS5405 | RS5405 | 0 | *Novosphingobium* | intra-specific assay |
| RS5405 | RS5405 | 0 | *Novosphingobium* | intra-specific assay |
| RS5405 | RS5405 | 0 | *Novosphingobium* | intra-specific assay |

**Supplementary Table5**
